# BIRD-Seq: B2 Protein Integrated End-to-End Pipeline for dsRNA Detection and Nanopore Sequencing for Virus Monitoring

**DOI:** 10.64898/2026.06.25.734422

**Authors:** Athanasios Nikolaos Xhurxhi, Subhankar Sahu, Hélène Scheer, Elliott F. Miott, Abdelmalek Alioua, Daniel Clesse, Baptiste Monsion, Julien Pompon, Sabine Szunerits, Todd Blevins, Christophe Ritzenthaler

**Affiliations:** Institut de Biologie Moléculaire des Plantes (IBMP), CNRS, Université de Strasbourg, Strasbourg, F-67084, France; MIVEGEC, Univ. Montpellier, IRD, CNRS, INRAE, 34394 Montpellier, France; UMR VIROLOGIE, École nationale vétérinaire d’Alfort, ANSES, INRAE, 94700 Maisons-Alfort, France; Danube Private University (DPU), Viktor-Kaplan-Straße 2, Geb. E, 2700 Wiener Neustadt, Austria; Univ. Lille, CNRS, Univ. Polytechnique Hauts-de-France, UMR 8520 - IEMN, F-59000 Lille, France

**Keywords:** double-stranded RNA (dsRNA), virus monitoring, sandwich assay, nanopore sequencing, infectious disease surveillance

## Abstract

Double-stranded RNA (dsRNA) is a near-universal hallmark of active viral infection. Despite its role as a pan-viral replication intermediate, dsRNA-centred technologies for virus monitoring remain scarce and largely rely on monoclonal antibody-based approaches that, while highly sensitive, are costly and difficult to engineer, scale, or integrate with downstream assays. Here, we present a modular and antibody-free pipeline for quantitative and qualitative dsRNA analysis built around an engineered B2 protein from Flock House virus with nanomolar-range binding affinity. The pipeline is compatible with absorbance- or luminescence-based measurement formats. In a sandwich assay configuration (Sand-BIRD), sub-ng mL^-1^ quantification of dsRNA is achieved, comparable to the gold standard J2 monoclonal antibody, directly from crude biological samples without RNA extraction. Sand-BIRD reliably detects viral infection in both plant samples (Tomato bushy stunt virus and Grapevine fanleaf virus) and mosquitoes (West Nile virus and Dengue virus) with commercial-grade reliability. dsRNA eluted from positive samples were further processed directly by Oxford Nanopore direct sequencing, enabling identification of virus species without prior sequence knowledge or total RNA extraction. Together, this work establishes an end-to-end, sequence-agnostic workflow for direct RNA-duplex quantification and sequencing (BIRD-Seq), which has compelling potential for emerging infectious disease surveillance and next-generation point-of-care (PoC) diagnostics.

**Technology Readiness:** BIRD-Seq is an integrated pipeline for agnostic dsRNA detection and sequencing designed for broad-spectrum virus monitoring. A Technology Readiness Level (TRL) 5 under NASA’s classification framework has been reached as BIRD-Seq has been validated in laboratory-relevant environments using real-world samples, including virus-infected plants and mosquitoes. The ELISA-based sensing platform employs engineered variants of the B2 protein (from Flock House virus) in a sandwich assay format for dsRNA capture and detection, achieving sensitivity comparable to the gold-standard J2 monoclonal antibody. Unlike traditional antibody-based methods, the B2 protein offers key practical advantages: straightforward production in bacterial expression systems, high versatility, and reduced manufacturing costs, as well as direct compatibility with crude extract monitoring, eliminating the need for RNA extraction. Captured B2/RNA duplexes can then be directly eluted from the ELISA microplates and subjected to downstream nanopore direct RNA sequencing, providing both quantitative and qualitative information on the underlying virus infection, a capacity enabled by the near-universal nature of dsRNA as a pathogen-associated molecular pattern. That said, further validation on large-scale field-collected and clinical samples will be essential before widespread deployment can be envisioned. While the B2 sandwich assay offers favorable cost-efficiency over antibody-based alternatives, the relatively high cost of Oxford Nanopore direct RNA sequencing remains an important economic constraint. Nevertheless, the growing importance of dsRNA detection across virus sensing, mRNA vaccine development, innate immunity research, and human disease diagnostics, combined with the increasing role of portable long-read sequencing in emerging infectious disease (EIDs) surveillance, positions BIRD-Seq as an innovative and competitive diagnostic platform.

**Highlights:**

– Double-stranded RNA (dsRNA) is one of the critical pathogen-associated molecular patterns for viral invasion in the host. A protein-based sandwich assay for dsRNA detection in crude biological samples with a sub-ng mL^-1^ order detection limit was developed, achieving similar sensing efficiency in comparison to expensive and proprietary monoclonal antibody-based ELISA methods.
– The quantitative detection of RNA duplex is coupled with an Oxford nanopore direct dsRNA sequencing method for virus species identification and qualitative analysis.
– This is one of the very first dsRNA-centered end-to-end workflows for virus monitoring and sequencing, validated for both infected plant and animal samples.

## Introduction

Emerging infectious disease (EID) events have risen significantly over the past decades globally, driven largely by socio-economic, environmental, and ecological changes [1,2]. The recent COVID-19 pandemic has made it clearer than ever that newly emerging viruses will pose an increasing worldwide threat in years to come, further intensified by factors like global mobility, urbanization, and climate change [2,3]. It also underscored certain limitations in current diagnostic frameworks and practical trade-offs: rapid and highly scalable diagnostics (such as RT-qPCR and antigen testing) tend to be targeted to specific pathogens, whereas broader, discovery-oriented platforms (like metagenomic sequencing) generally entail higher costs and scalability challenges [4,5]. To tackle the adverse impact of the viral diseases on human health, crop production, and food safety, there remains a need for adaptable, rapid, and cost-effective methodologies capable of not only probing generic viral infection but also identifying novel viruses [5–7], as by definition, EIDs cannot be detected by targeted approaches that require prior knowledge of the pathogens. Owing to its near-universal presence during viral replication of both RNA and DNA viruses and its inherent thermodynamic stability, double-stranded RNA (dsRNA) is increasingly recognized as a highly reliable biomarker for the agnostic detection of viruses, offering a means to identify emerging pathogens without relying on prior knowledge of the given virus [8–11]. Surprisingly, a widespread application of dsRNA-based approaches for virus detection remains largely underexplored, with existing work focusing predominantly on either dsRNA detection [8–10] or RNA sequencing in isolation [12–14], but rarely integrating both approaches, which is a central objective of the present study.

Double-stranded RNA serves as a highly versatile biological scaffold that mediates diverse cellular functions [15]. While RNA duplex motifs naturally occur in various cellular transcripts, including tRNAs, rRNAs, long non-coding RNAs, and mRNA untranslated regions (UTRs), long dsRNA species are tightly regulated in most species. Long RNA duplexes are widely recognized as pathogen-associated molecular patterns (PAMPs) that rapidly accumulate during viral infection and replication [16,17]. Consequently, multicellular organisms have evolved sophisticated innate immune responses to detect and process these viral dsRNAs (V-dsRNAs), utilizing intracellular sensors such as protein kinase R (PKR), Toll-like receptors (TLRs), retinoic acid-inducible gene I (RIG-I), and melanoma differentiation-associated protein 5 (MDA5) in vertebrates [18,19], as well as Dicer enzymes in plants [20]. Historically, this has established dsRNA primarily as a virological “danger” molecule that alerts the host immune system to actively replicating viruses [15]. Beyond infectious disease diagnostics, dsRNA is also increasingly recognized as a multifaceted molecule across broader biomedical fields. It is notably a major contaminant requiring stringent clearance during mRNA vaccine production [21], and its aberrant accumulation is increasingly linked to the pathogenesis of numerous neurodegenerative, autoimmune, and inflammatory human diseases [22–25].

In the context of broad virus surveillance, metagenomic next-generation sequencing (mNGS) and metatranscriptomics have been recognized as effective, sequence-independent avenues for pathogen monitoring. These approaches enable the unbiased detection of known and novel viral agents across a diverse range of clinical, environmental, and ecological samples [26,27]. Their implementation is often constrained by the overwhelming burden of host nucleic acids in most biological matrices. Viral sequences may represent less than 0.1% of total sequence reads and results in shallow viral coverage and reduced sensitivity [28]. To circumvent this limitation, techniques such as VANA (virion-associated nucleic acids) [29] have been implemented to increase the relative proportion of viral reads prior to library preparation. VANA relies on the physical purification of virions to enrich encapsulated viral genomes before shotgun sequencing. Although VANA-based metagenomics has contributed to the discovery of more than 75 novel virus species over the past decade [30], it is inherently confined to viruses that produce stable virion particles, thereby overlooking capsidless or persistently replicating agents. Conversely, targeted enrichment of dsRNA can serve as a complementary and biologically informative alternative. Because dsRNA is an obligate replication intermediate for nearly all RNA viruses yet largely absent in uninfected eukaryotic cells [15], its isolation and sequencing can provide direct evidence of active viral replication and facilitate the identification of *bona fide* viral agents.

Several well-established methodologies are currently in use for the isolation and detection of dsRNA. Traditional cellulose-based chromatography has long served as a standard procedure for purifying viral dsRNA and reliably capturing high-molecular-weight RNA duplexes [31]. More recently, commercial kits utilizing the *Arabidopsis* DRB4 protein (e.g., Plant Viral dsRNA Enrichment Kit; MBL Life Science) have streamlined this process, providing user-friendly workflows for dsRNA enrichment [32]. In the 1990s, Schönborn et al. developed a panel of four monoclonal antibodies (J2, J5, K1, K2) against dsRNA, among which J2 has become the gold standard for direct visualization and diagnostic detection [33]. Initially reported to bind RNA duplexes of at least 11 bp in a sequence-independent manner, J2 was then applied to sandwich ELISA and immunoblotting for plant virus detection before being adapted for immunofluorescence imaging in animal cells, where it demonstrated that positive-strand RNA and DNA viruses, but not negative-strand RNA viruses, produce detectable dsRNA during infection [33–35]. J2-based immunoprecipitation has since been further coupled to next-generation sequencing for transcriptome-wide dsRNA profiling [36]. Subsequent biochemical and structural studies have, however, revealed important limitations. Atomic force microscopy showed a minimum binding threshold of approximately 40 bp and identified a preferential affinity for AU-rich sequences [37], while the recently resolved co-crystal structure of J2 with dsRNA rationalizes both observations: J2 tracks the minor groove via a staggered 8-bp footprint, with robust binding requiring at least 14 bp, and GC-rich duplexes are disfavored due to their narrower minor groove geometry [38]. Beyond these binding biases, J2 is costly to produce, frequently proprietary, and difficult to engineer into reporter fusions. Finally, a limitation shared by nearly all dsRNA-sensing methodologies is the reliance on prior organic solvent-based total RNA extraction, which represents a key bottleneck to high-throughput application.

To overcome these methodological and economic limitations, recent advances have shifted toward utilizing dsRNA-binding proteins (dRBPs) as biological dsRNA detection scaffolds. One such dRBP is the B2 protein from Flock House virus (FHV), which possesses high affinity for RNA duplexes. Functioning naturally as a viral suppressor of RNA interference (RNAi), the B2 homodimer forms a four-helix bundle that interacts specifically with the ribose-phosphate backbone of A-form RNA duplexes in a sequence-independent manner [39,40]. Previously, Ritzenthaler et al. demonstrated that B2 is a highly efficient reporter for dsRNA binding *in vitro* using approaches such as northwestern blotting and fluorescence labelling, as well as *in vivo* via fluorescence microscopy [9]. Beyond simple detection, the binding properties of B2 have been leveraged to immunocapture dsRNA-bound proteins, yielding valuable insights into viral replication complexes [41]. Specifically, a B2-green fluorescent protein fusion (B2:GFP) was utilized to pull down dsRNA and its associated proteins from plant extracts. Furthermore, for viral ecology studies, this B2-based method has proven highly cost-effective for dsRNA purification from total RNA pools prior to high-throughput sequencing (HTS), showcasing a per-reaction cost of just $4.47 compared to $35.34 for DRB4-based approaches [42]. In parallel, the B2 scaffold has been applied to detect dsRNA in *in vitro*-transcribed (IVT) mRNA samples using Biolayer Interferometry (BLI) [43]. Recently, two other dsRBD-based detection approaches have been described, utilizing a family of adenosine deaminases acting on RNA (ADAR) and PKR domains for sensitive dsRNA visualization and quantification [44,45].

Comprehensive sequence-based characterization for taxonomic classification and epidemiological tracking has been transformed in recent years by portable, long-read direct sequencing technologies, which hold significant potential for on-field pathogen monitoring [46,47]. Oxford Nanopore Technologies (ONT), in particular, has rapidly gained traction as a transformative platform for pathogen surveillance and viral metagenomics [48,49]. ONT offers distinct advantages over traditional short-read HTS approaches, including rapid sample-to-identification turnaround times, often under 6 hours via real-time sequence acquisition, and the ability to produce long reads that facilitate *de novo* genome assembly and structural variant monitoring without fragmentation artifacts [50]. Combining dsRNA enrichment strategies with nanopore sequencing enhances viral RNA detection sensitivity while simultaneously depleting abundant host rRNA and mRNA transcripts. Recently, a comparative study in grapevine viral diagnostics utilizing a ‘NanoViromics’ approach showed that dsRNA-based nanopore libraries revealed a much larger proportion of viral reads (∼85–95%) than rRNA-depleted total RNA libraries (about 6–21%), and dsRNA-based sequencing identified more lower-abundance viruses [13]. Furthermore, nanopore sequencing achieves unbiased viral discovery across taxonomically diverse lineages due to its amplification-free nature [51]. Additionally, the direct RNA sequencing platform enables the detection of post-transcriptional modifications (e.g., pseudouridine, N6-methyladenosine), preserves strand-specific information, and supports real-time sequence analysis, facilitating the dynamic decision-making critical for outbreak investigations [52,53]. For example, genomic surveillance in the Ebola epidemic in West Africa was one of the very early large-scale portable applications of the MinION sequencer [54]. The usage is further steadily increasing for both animal virology, such as dengue [55], Zika [56], SARS-CoV-2 [57], etc., and plant virology, including plum pox virus [58], cassava mosaic disease [48], viruses in grapevines [13], etc.

Leveraging the collective advantages of nanopore sequencing and the highly sensitive RNA duplex-binding properties of the B2 protein, we developed an integrated, extraction-free pipeline for dsRNA sensing and subsequent sequencing, designated as **BIRD-Seq** (B2-Integrated dsRNA detection and sequencing) (**Figure 1**). By exploring the structural adaptability of the B2, we engineered versatile reporter fusions (NanoLuc luciferase and alkaline phosphatase) to construct a solid-phase sandwich assay (**Sand-BIRD**) capable of directly quantifying viral dsRNA in crude plant and mosquito extracts. The assay can selectively detect dsRNA of natural viral origin, reaching an LOD and LOQ of around 0.7 and 2.4 ng mL^-1^, respectively. This platform can rapidly screen complex matrices without any prior RNA purification and was validated for dsRNA-mediated pathogen sensing across a diverse panel of viruses, including Tomato bushy stunt virus (*Tombusvirus lycopersici*, TBSV), Grapevine fanleaf virus (*Nepovirus foliumflabelli*, GFLV), West Nile virus (*Orthoflavivirus nilense*, WNV), and Dengue virus (*Orthoflavivirus denguei*, DENV). Following positive detection and quantification via Sand-BIRD, dsRNA can be directly eluted from the B2-bound surface and coupled to a downstream nanopore direct RNA sequencing protocol (**dsNanoSeq**) for qualitative analysis, enabling accurate mapping to viral genome repositories to identify the exact viral agent present. This integrated framework represents one of the first dsRNA-focused virus surveillance platforms capable of distinguishing actively infected samples by quantifying dsRNA levels directly in unpurified, crude extracts. Crucially, this process simultaneously acts as an enrichment step to capture and sequence predominantly virus-derived dsRNA species, providing a streamlined, scalable, end-to-end solution for agnostic pathogen discovery.

**Figure 1.**
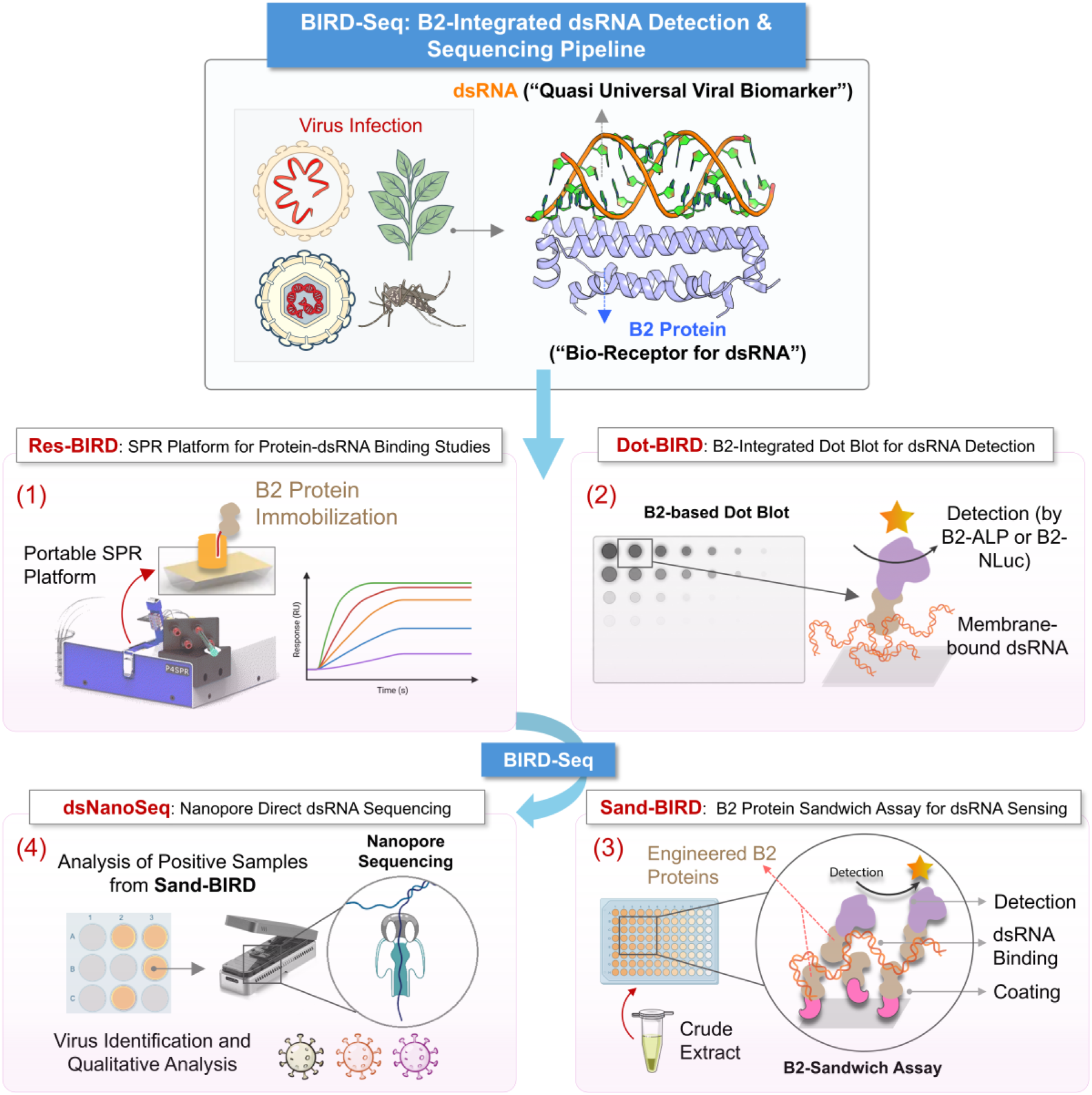
Schematic overview of the B2 protein-based end-to-end agnostic workflow for dsRNA detection and virus monitoring. Engineered variants of the B2 protein (from Flock House virus; 1-73 amino acids, PDB ID: 2AZ0) were exploited as a highly efficient bioreceptor for dsRNA detection (a quasi-universal viral biomarker) and were integrated across various analytical platforms to monitor dsRNA *in vitro*, in crude extracts, and during viral infection in plants and insects. The pipeline is divided into four distinct functional modules: (1) **Res-BIRD**, a semi-portable surface plasmon resonance (SPR) platform utilized for characterizing protein-dsRNA binding kinetics; (2) **Dot-BIRD**, a B2-integrated dot blot assay for the rapid detection of membrane-bound dsRNA; (3) **Sand-BIRD**, B2 protein integrated sandwich assay for dsRNA monitoring directly in crude samples (employing engineered B2 proteins such as B2-mScarlet for dsRNA capture alongside B2-ALP or B2-NLuc for enzymatic revelation); and (4) **dsNanoSeq**, a nanopore direct dsRNA sequencing methodology for downstream viral species identification and qualitative analysis of positive samples. Collectively, these platforms constitute the **BIRD-Seq** (B2-Integrated dsRNA Detection & Sequencing) pipeline, providing a comprehensive RNA extraction-free framework for dsRNA detection and virus identification.

## Results

### Surface-immobilized B2 is compatible for dsRNA binding with an affinity comparable to the gold standard J2 mAb

B2 protein (from FHV) is an efficient dsRNA bioreceptor, and its binding affinity to short dsRNA and long 700 bp unmodified dsRNA has been characterized earlier by electrophoretic mobility shift assay (EMSA) [39] and biolayer interferometry (BLI) [43], respectively. EMSA studies reported a binding affinity (*K*_D_) of 1.4 nM for B2 fused to a maltose binding protein (MBP-B2) toward a 19-bp RNA duplex bearing 5′-phosphate and 3′ 2-nt overhang. The latter study reported a *K*_D_ ∼60 nM for B2 (carrying a smaller AviTag) toward unmodified 700 bp dsRNA; binding was further weakened for dsRNA bearing uridine modifications such as pseudouridine (ψ), N1-methylpseudouridine (m1ψ), and 5-methoxyuridine (5moU) — with *K*_D_ values ranging from 405 to 948 nM.

In this work, the binding affinity of B2 for long *bona fide* viral dsRNA (V-dsRNA) was determined by a portable surface plasmon resonance (SPR) setup. The first step was to establish an oriented immobilization strategy for B2 on a gold-coated SPR chip. For this, the SpyTag003/SpyCatcher003 strategy (referred to as SpT and SpC) [59], wherein the SpT part spontaneously forms a covalent bond *in vitro* with a SpC partner (**Figure S1C, S2A),** was employed to tether the B2 [60]. As an initial step, a Cys-SpyCatcher003 protein (C-SpC) containing cysteine positioned at the N-terminal side of the construct (following the N-terminal 6× His tag, separated by a short Gly/Ser linker from the core SpyCatcher003 domain; **Table S1**) was immobilized on the gold surface (**Figure 2A**, Step-1). Since SpC naturally lacks cysteine (Cys) residues, one was introduced at the N-terminus side, enabling direct immobilization of SpC on the gold surface via thiol (-SH) chemistry. Addition of a B2 protein fused to a SpyTag003 (B2-SpT) (**Figure 2A**, Step-2) onto this surface resulted in an increase in the SPR response units (RU) to about ∼125 RU after the washing/dissociation step (**Figure S2B**), indicating successful B2 immobilization on the surface with a coverage of ∼0.12 ng mm^-2^ (assuming that 1 RU approximately corresponds to 1 pg mm^-2^ of protein [61]. Addition of V-dsRNA, commercially known as ‘Larifan’ (**Figure 2A**, step 3), further resulted in an increase in RU signal consistent with binding to the B2-coated surface (**Figure S2C**). In contrast, poly U single-stranded RNA (ssRNA), even at a 50-fold higher concentration (**Figure S2D**), showed a negligible response. These results confirm the selectivity of the B2 bioreceptor and validate that the surface-anchored B2 is functional for selective dsRNA detection.

**Figure 2.**
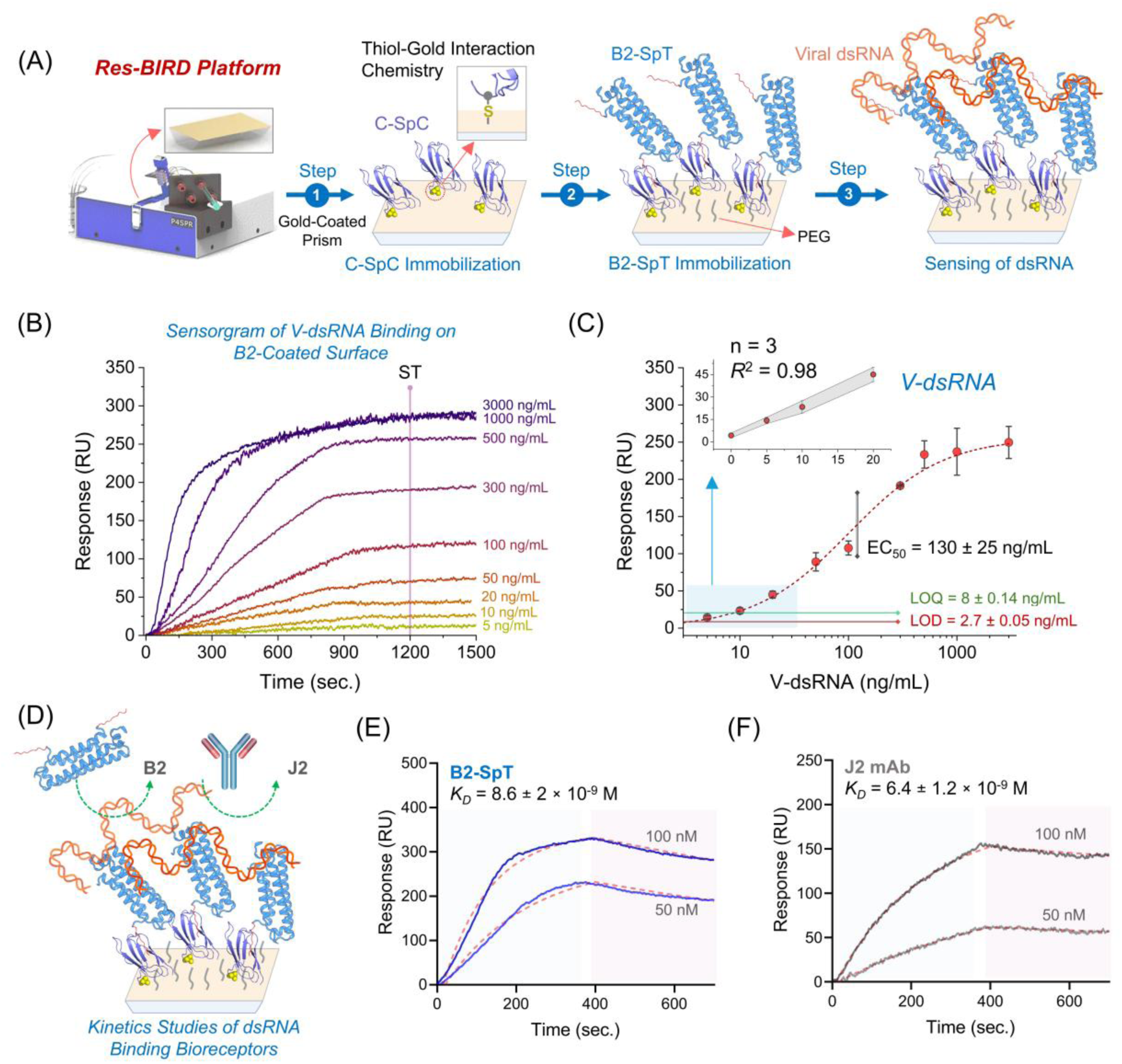
B2/dsRNA binding kinetics studies by surface plasmon resonance (SPR). **(A)** Schematic representation of the SPR platform employed for B2/dsRNA binding kinetics studies. The portable P4SPR from Affinité Instruments is shown along with the pristine gold SPR prism. The B2 bioreceptor was immobilized in an oriented manner onto the gold-coated prism by the cysteine SpyCatcher003 (C-SpC) and SpyTag003 (SpT) interaction chemistry. **Step-1:** C-SpC is immobilized onto the gold surface through the Cys functional moiety (highlighted by yellow color ball-shaped atoms). **Step-2:** The surface is passivated with an antifouling PEG layer, and B2-SpyTag003 (B2-SpT) is attached subsequently to the surface, acting as a molecular carpet for dsRNA binding/capturing. In the schematic, the SpT domain (red) is shown covalently coupling to the surface-bound C-SpC. **Step-3:** *bona fide* viral dsRNA (V-dsRNA) (shown in orange) is specifically recognized and captured by the B2 bioreceptor. **(B)** SPR response profiles of V-dsRNA binding by B2 protein across an increasing nucleic acid concentration of 5 – 3000 ng mL^-1^ (ST: represents the steady state time point). **(C)** The steady state RU response (obtained from Figure 2B) is shown as a function of increasing V-dsRNA concentration (ranging across 5-3,000 ng mL^-1^). A 4-parameter logistic (4PL) curve fitting reveals the sigmoidal trend with EC_50_ value = 130 ± 25 ng mL^-1^. The lower region of the standard curve is given as inset (the shadowed region represents the standard deviation associated with each measurement). Each data point represents mean ± standard error of the mean (SEM), n = 3 technical replicates. **(D)** The surface captured V-dsRNA (in Figure 2A, step-3) serves as a uniform saturated bio-layer for studying the kinetics of RNA-duplex binding bioreceptors (such as B2 protein or J2 mAb). Schematic of the SPR prism showing B2 protein and J2 mAb binding kinetics studies with the V-dsRNA nucleic acid species. SPR sensorgrams and binding affinity determination of **(E)** B2 protein and **(F)** J2 mAb for *bona fide* viral dsRNA. A dissociation constant (*K*_D_) of 8.6 nM and 6.4 nM was obtained for B2 and J2, respectively.

In a concentration-dependent titration of V-dsRNA, the SPR signal increased proportionally with dsRNA captured by B2, reaching saturation at approximately 1000 ng mL^-1^ (F**igure 2B**). Steady state analysis (SSA) of this titration data yielded a half maximal effective concentration (EC_50_), limit of detection (LoD), and limit of quantification (LoQ) of 130 ± 25, 2.7 ± 0.05, and 8 ± 0.14 ng mL^-1^, respectively (**Figure 2C**) for pure V-dsRNA (in 1× PBS-T buffer). The strong avidity of B2 nearly eliminates dissociation (**Figure 2B**), creating a uniform dsRNA layer (with a stable, consistent baseline). In principle, this is a surface saturated with dsRNA and can be applied for measuring the affinity of a dsRNA-binding bioreceptor. Here, each bioreceptor binding site to the surface immobilized dsRNA can be treated as independent from one another; therefore, the 1:1 binding model can be applied for measuring the affinity. A similar strategy was also substantiated earlier within a BLI experimental setup by Silas et al. [43]. Our SPR experiment setup revealed a lower nanomolar order affinity of B2 for *bona fide* dsRNA, with an apparent *K*_D_ of 8.6 ± 2 nM (**Figure 2E**). When J2 mAb was titrated over the same V-dsRNA surface, it yielded an apparent *K*_D_ of 6.4 ± 1.2 nM, comparable to that of the B2 bioreceptor (**Figure 2F**). This platform, referred to here as B2 integrated SPR (**Res-BIRD**), not only validates B2 as an alternative dsRNA-sensing bioreceptor but also confirms that B2 retains its dsRNA-binding properties in a surface-anchored format.

### B2 can be structurally engineered to incorporate reporter proteins for dsRNA monitoring

Having demonstrated that the B2 bioreceptor exhibits high affinity for V-dsRNA, we then assessed whether this core binding domain could be modularly fused to functional reporter units without compromising target recognition. To this end, we generated a series of C-terminal fusion constructs (**Figure 3A**). These included a fluorescent module (B2-mScarlet) (**Figure 3A, S4**), intended primarily for surface capture, alongside two distinct enzymatic reporters for downstream signal generation. Specifically, we incorporated bacterial alkaline phosphatase (B2-ALP) (**Figure 3A, S5**), a dimeric enzyme that produces a colorimetric readout via dephosphorylation of *p*-nitrophenyl phosphate (pNPP) [62,63], and NanoLuc luciferase (B2-NLuc) (**Figure 3A, S6**), a monomeric reporter that generates a bioluminescent signal through the oxidation of a furimazine substrate [64]. Sequence details of all engineered B2 constructs are provided in **Table S1**.

**Figure 3.**
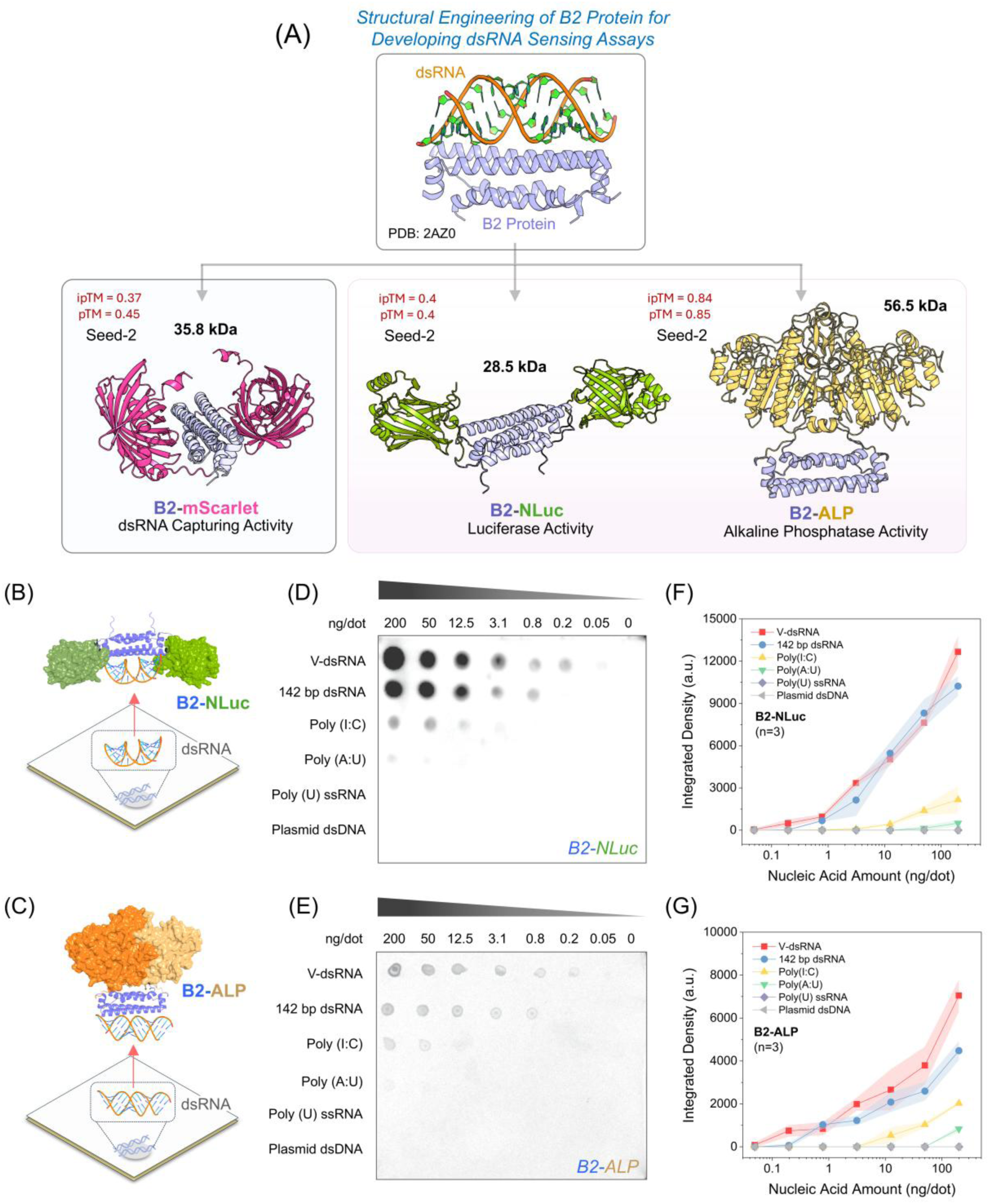
Structural engineering of B2 fusion proteins and their application for dsRNA detection in a dot-blot format (Dot-BIRD assay). **(A)** The dsRNA binding domain of B2 protein (1-73 amino acids, PDB ID: 2AZ0) is structurally engineered to fuse with functional reporter modules, and distinct dsRNA bioreceptors are created: B2-mScarlet (utilized for surface coating and dsRNA capture), alongside B2-NanoLuciferase (B2-NLuc) and B2-Alkaline Phosphatase (B2-ALP) (utilized for downstream enzymatic revelation of captured dsRNA). The AlphaFold-3 generated models are shown for B2-mScarlet, B2-NLuc, and B2-ALP. Respective pTM (predicted Template Modeling score) and ipTM (interface predicted Template Modeling) scores are shown beside each predicted structure, along with the molecular weight (M_W_). Schematic diagram of membrane-bound dsRNA detection by **(B)** B2-NLuc as an engineered scaffold using chemiluminescence and **(C)** B2-ALP employing colorimetric detection. Representative dot blot membrane images (Dot-BIRD assay) of the **(D)** B2-NLuc and **(E)** B2-ALP construct. The blots show an array of different nucleic acid species (viral dsRNA of natural origin (V-dsRNA), 142 bp dsRNA standard, Poly(I:C), Poly(A:U), Poly(U) ssRNA, and dsDNA) tested across a range of serial fourfold dilution concentrations (0-200 ng per dot). The readouts demonstrate that while both B2 scaffolds recognize all tested RNA duplex species in a concentration-dependent manner, they possess a significantly higher binding affinity for *bona fide* dsRNA targets (such as V-dsRNA and 142 bp dsRNA) compared to the synthetic dsRNA moieties and reveal no response for ssRNA or dsDNA. Quantitative integrated density plots derived from dot blot (using ImageJ) for the **(F)** B2-NLuc and **(G)** B2-ALP variants. The engineered B2 biosensors strictly retain their structural preference, with no cross-reactivity observed against single-stranded RNA (like Poly(U)) or plasmid double-stranded DNA (dsDNA), even at the highest tested concentrations. Data are presented as the mean, and the shadowed region associated with each trendline represents the standard deviation (n = 3) involved with each measurement.

Because fusing a small binding domain to relatively bulky heterologous tags can introduce steric hindrance and impair proper folding, we first purified each variant (**Figure S7**) and then validated their dsRNA-binding affinity using the Res-BIRD platform (**Figure S8**). By flowing the purified fusion proteins over a surface saturated with V-dsRNA, we monitored their association and dissociation kinetics. The SPR analysis confirmed that all three engineered B2 constructs retained their ability to recognize and bind RNA duplexes. While the addition of the reporter tags resulted in a reduction in binding strength compared to the wild-type protein, as expected, the fusions still maintained robust affinities in the nanomolar range with *K*_D_ of 28 ± 2 nM for B2-mScarlet, 54 ± 14 nM for B2-NLuc, and 105 ± 16 nM for B2-ALP, respectively (**Table S2**).

### B2 is highly efficient for dsRNA detection in a conventional dot blot format

After confirming nanomolar binding affinity for the engineered B2 constructs using Res-BIRD, we adapted the traditional nucleic acid dot blot assay into a direct B2-based format for dsRNA detection, termed **Dot-BIRD** (B2-integrated dsRNA detection dot blot assay). Using B2-NLuc (**Figure 3B**) and B2-ALP (**Figure 3C**) recombinant proteins as bioreceptors, we explored whether they could generate a functional signal directly proportional to the amount of target RNA duplex immobilized on a membrane. Serial fourfold dilutions of various nucleic acid species (ranging from 200 to 0.05 ng/dot) were spotted to obtain binding profiles. For the B2-NLuc bioreceptor, the addition of the furimazine substrate yielded a bioluminescent signal proportional to the amount of immobilized natural-borne V-dsRNA and 142 bp dsRNA standard (**Figure 3D**). A similar dose-response profile was obtained using the B2-ALP variant with its BCIP/NBT colorimetric substrate (**Figure 3E**), demonstrating that B2 is functional for dsRNA detection regardless of the fused reporter tag. The complete Dot-BIRD workflow is illustrated in **Figure S9**. To quantify these visual dot blot readouts, densitometric analysis was performed using ImageJ, yielding integrated density plots that revealed concentration-dependent responses (**Figure 3F, G**).

Evaluation of target selectivity revealed a binding preference of the B2 domain for *bona fide* natural dsRNA. While both B2-NLuc and B2-ALP moderately detected synthetic RNA duplex mimics such as Poly(I:C) and Poly(A:U), binding to *bona fide* natural dsRNA targets was markedly favoured. This preference was corroborated by our Res-BIRD experiments, in which testing B2-NLuc and B2-ALP against immobilized synthetic dsRNA species yielded at least a two-fold increase in the *K*_D_, indicating reduced affinity (**Figure S10**). Furthermore, we observed no cross-reactivity with off-target nucleic acids, such as ssRNA (Poly(U)) or dsDNA, even at the maximal concentration (200 ng per dot) tested. Overall, the Dot-BIRD setup demonstrates that engineered B2 proteins remain highly functional in a membrane-bound matrix and can be effectively implemented in standard solid-phase assays to detect dsRNA species. By achieving a limit of detection (LOD) of ∼0.2 ng/dot for dsRNA of natural viral origin, the assay matches the sensitivity of traditional antibody-based techniques [81] while offering the distinct advantages of a single-step, single-molecule sensor system.

### B2 protein integrated sandwich assay (Sand-BIRD) is a highly efficient method for dsRNA detection

Transitioning from membrane-based detection, we designed a B2 protein integrated sandwich assay adapted to 96-well microplate (**Sand-BIRD**) inspired by traditional double-antibody sandwich ELISAs (**Figure 4A**). In this setup, B2-mScarlet serves as the solid-phase bioreceptor to capture dsRNA. To maximize the platform’s versatility, we evaluated two distinct reporter systems for the revelation of surface-bound RNA duplexes: a colorimetric approach utilizing the dimeric B2-ALP, and a luminescent approach using the monomeric B2-NLuc. Although antibody-based ELISA have been widely adapted for dsRNA detection using colorimetric, fluorescent, or chemiluminescent readouts [65], we are not aware of a prior non-antibody-based sandwich assay specifically designed for dsRNA detection.

**Figure 4.**
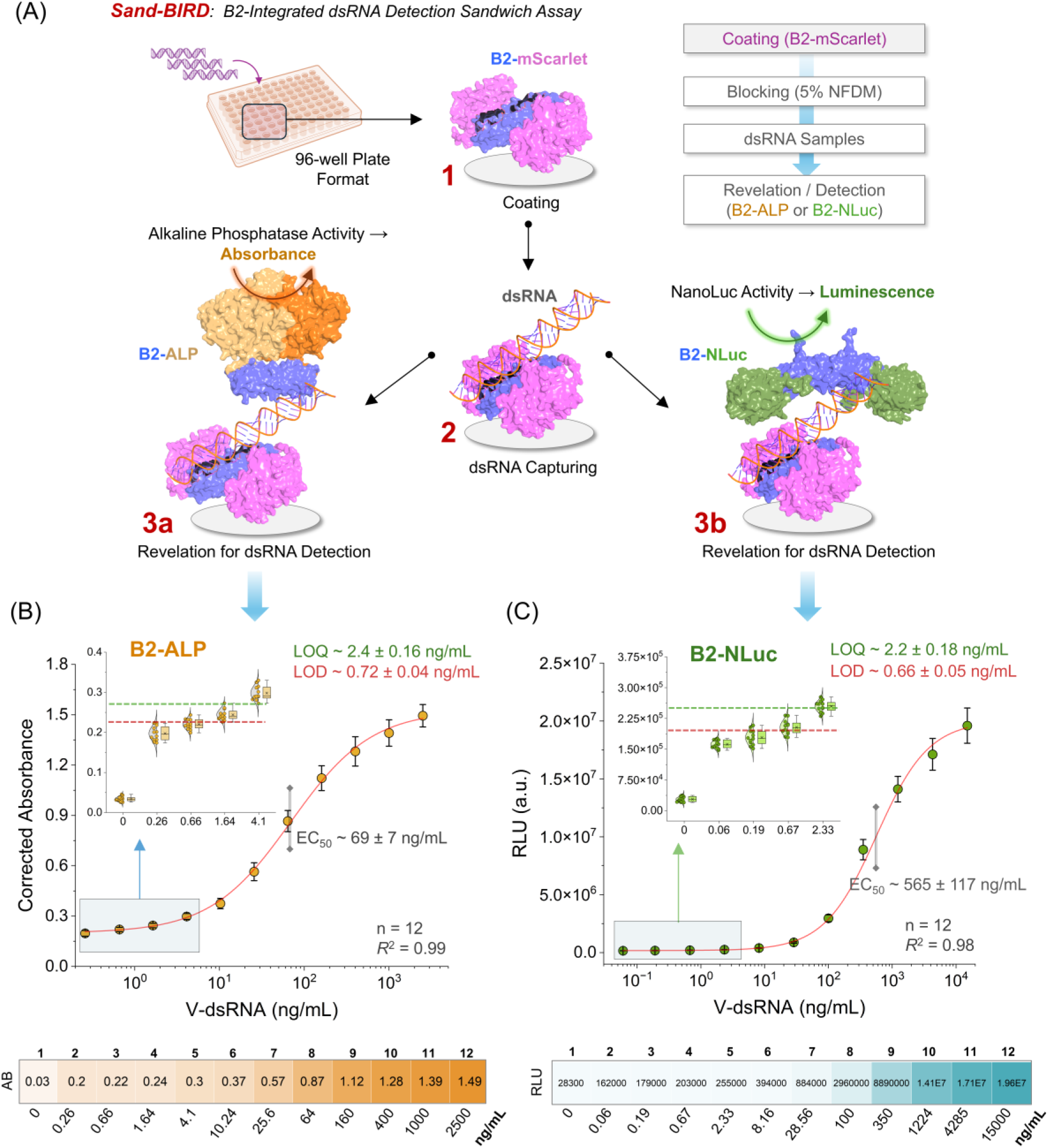
Development and analytical performance of the B2-integrated dsRNA detection sandwich assay (Sand-BIRD). **(A)** Schematic pipeline of the Sand-BIRD assay adapted for a high-throughput 96-well microplate format. The assay utilizes the engineered B2-mScarlet fusion protein for surface coating (1) to efficiently capture the target viral double-stranded RNA (V-dsRNA) (2). Following the capture step, enzymatic revelation is performed utilizing either the B2-Alkaline Phosphatase (B2-ALP) variant for a colorimetric readout or the B2-NanoLuciferase (B2-NLuc) variant for a luminescent readout (3a/3b). **(B)** Standard calibration curve and performance metrics for the colorimetric B2-ALP detection module. The assay utilizes *p*-nitrophenyl phosphate (pNPP) as a substrate. The y-axis represents corrected absorbance, representing the absorbance measured at 405 nm (with background correction at 492 nm). This setup achieves a limit of detection (LOD) of 0.72 ng mL^-1^, a limit of quantification (LOQ) of 2.4 ng mL^-1^, and a half-maximal effective concentration (EC_50_) of 69 ng mL^-1^. The bottom panel shows the mean corrected absorbance values of each tested V-dsRNA concentration. **(C)** Standard calibration curve and performance metrics for the luminescent B2-NLuc detection module. The assay employs a furimazine substrate to produce a luminescence readout, achieving an LOD = 0.66 ng mL^-1^, an LOQ = 2.2 ng mL^-1^, and an EC50 = 565 ng mL^-1^. The y-axis shows photon count in terms of relative light units (RLU). The bottom panel shows the mean RLU values of each tested V-dsRNA concentration. Both Sand-BIRD configurations (B2-ALP and B2-NLuc) demonstrate a broad dynamic range with very high sensitivity for the quantitative detection of *bona fide* viral dsRNA targets. Data are presented as mean ± SD (n = 12). The figure inset in (B and C) shows raw data points of the lower asymptote region of the standard curve as a raincloud plot, where the box plot indicates the median and interquartile range, the solid scatter points represent individual replicate measurements, and the shaded region beside the box plot represents the data distribution density.

When tested against *bona fide* V-dsRNA in buffer (1× PBS), both reporter systems exhibited very high sensitivity, achieving sub-nanogram-per-millilitre order limits of detection. Specifically, B2-ALP and B2-NLuc reached LODs of 0.72 ± 0.04 ng mL⁻¹ and 0.66 ± 0.05 ng mL⁻¹, respectively (**Figure 4B, C**). The limit of quantification (LOQ) and half-maximal effective concentration (EC50) for the B2-ALP scaffold were 2.4 ± 0.16 ng mL⁻¹ and 69 ± 7 ng mL⁻¹, while the B2-NLuc system yielded an LOQ of 2.2 ± 0.18 ng mL⁻¹ and an EC_50_ of 565 ± 117 ng mL⁻¹. The assays demonstrated a broad dynamic range, ranging from 2.4 to 400 ng mL⁻¹ for ALP and 2.2 to 1,200 ng mL⁻¹ for NLuc, validating their robust quantitative capacity across a large range of dsRNA concentrations (**Figure S11A-B**). Importantly, the Sand-BIRD platform preserved the stringent specificity seen in our dot blot assays, showing no measurable response to off-target ssRNA or dsDNA. The signal was also essentially unaffected when V-dsRNA was tested in a complex simulated mixture containing ssRNA and dsDNA, supporting the assay’s robustness for RNA duplex quantification in heterogeneous nucleic acid samples (**Figure S11C**).

### Sand-BIRD detects dsRNA in crude samples from virus-infected plants and insects

A common limitation of antibody-based methods for dsRNA detection is the requirement for prior total RNA extraction [11]. Direct detection of RNA duplexes in crude biological samples remains challenging due to matrix complexity, including in plant tissues, where high levels of polyphenols, polysaccharides, and other inhibitory molecules can interfere with the analysis [66]. To address this issue, the Sand-BIRD assay was designed to be directly compatible with crude extracts analysis. The sample preparation workflow involves simple tissue grinding in an extraction buffer, followed by a brief clarification step at 14,000 × *g* for 5 minutes (**Figure 5A**). The resulting supernatant was then transferred directly to the assay plate, enabling a scalable RNA-extraction-free approach. We privileged the B2-mScarlet/B2-ALP over the mScarlet/B2-nLuc configuration for detection, owing to the simple and robust colorimetric readout offered by ALP. Furthermore, since most commercial plant virus ELISA kits use ALP as the reporter enzyme - rather than horseradish peroxidase, which is often unsuitable due to high peroxidase activity in plant tissue - ALP allowed the Sand-BIRD assay to be directly benchmarked against established commercial virus detection kits.

**Figure 5.**
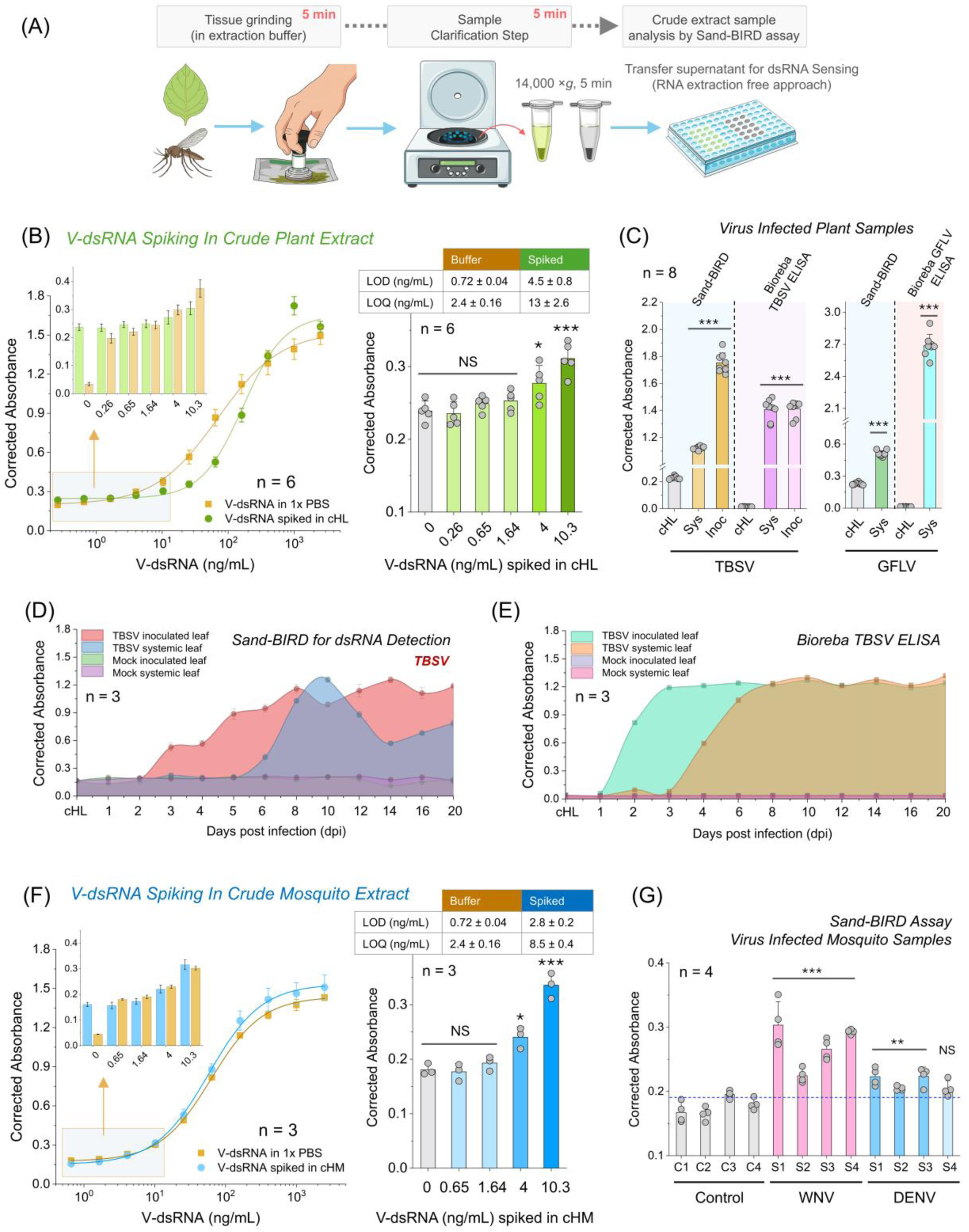
Validation of the B2 protein sandwich assay (Sand-BIRD) for viral dsRNA monitoring in complex biological samples. **(A)** Schematic representation of the rapid, RNA-extraction-free sample preparation workflow for Sand-BIRD. The pipeline utilizes direct mechanical cell lysis and clarification of complex biological matrices, such as infected plant tissues or mosquito vectors, to release dsRNA for downstream Sand-BIRD analysis. **(B)** Sand-BIRD assay (using B2-ALP) standard calibration curve comparing the response for V-dsRNA detection in 1× PBS (orange) and when spiked in crude plant extract (1:10 w/v tissue dilution; green) to validate biosensor performance in a complex matrix. cHL represents control healthy leaves. The lower concentration region of the signal comparison is provided in the inset as a bar plot. The lower concentration range (0-10.5 ng mL^-1^) response of the V-dsRNA spiked in healthy plant extract is also shown (right) as a scatter-bar plot along with the comparison table for LOD/LOQ figure of merits. **(C)** Comparative analysis of Tomato bushy stunt virus (TBSV) infection in crude *N. benthamiana* plant extracts. The dsRNA levels in inoculated (Inoc) and systemic (Sys) leaves, compared to control healthy leaves (cHL), are plotted alongside the identical samples tested with a commercial TBSV virus detection ELISA (from Bioreba), demonstrating the correlation between the dsRNA presence and the viral load. Comparative analysis of Grapevine fanleaf virus (GFLV) infection is also shown beside. Similar to the TBSV model, the Sand-BIRD assay successfully detected dsRNA levels in infected versus healthy plant extracts. The assay is compared against a commercial GFLV ELISA to compare universal dsRNA biomarker levels with specific viral load. **(D)** Time-course assay tracking the progression of TBSV infection in *N. benthamiana*. The Sand-BIRD platform monitors how the levels of viral dsRNA accumulate and propagate in both inoculated (red) and systemic leaves (blue) over the different days post-infection (dpi). Response profiles of the mock samples were shown in each case as control samples. **(E)** Time-course tracking of the identical TBSV-infected plant samples tested in parallel with the commercial TBSV ELISA. The plot confirms the infection dynamics by monitoring the increase in viral load over time, which correlates well with the dsRNA level from the Sand-BIRD (Figure 5C). **(F)** Standard calibration curve evaluating the Sand-BIRD assay performance for V-dsRNA spiked into crude healthy mosquito extracts (1:10 w/v tissue dilution; blue) and in 1× PBS (orange), further revealing the quantitative capability of the biosensor in animal vector matrices. cHM denotes control healthy mosquitoes. The signal comparison of the lower concentration region is provided in the inset as a bar plot. The lower concentration range (0-10.5 ng mL^-1^) response of the V-dsRNA spiked in healthy mosquito extract is also shown (right) as a scatter-bar plot along with the comparison table of LOD/LOQ figure of merits. **(G)** Validation of the Sand-BIRD platform in virus-infected mosquito samples (*Ae. aegypti*). The plot compares dsRNA levels measured in healthy control mosquitoes (cHM) against those infected with West Nile Virus (WNV) and Dengue Virus (DENV). Data are presented as mean ± SD, and the number of data points (n values) is given for each case inside the plot. The *p*-value was calculated by Welch’s two-sample t-test, and it is indicated on the bar charts (ns: *p* > 0.05, *: 0.05 > *p* > 0.01, **: 0.01 > *p* > 0.001, and ***: *p* < 0.001).

To rigorously quantify matrix interference caused by cellular components, we compared standard calibration curves of *bona fide* V-dsRNA in buffer alone with V-dsRNA spiked into crude matrices. First, to mimic an infected plant environment, we spiked a known quantity of V-dsRNA into a crude extract derived from control healthy leaves (cHL) of *Nicotiana benthamiana*. While the standard curve in pure buffer yielded a LOD of 0.72 ± 0.04 ng mL⁻¹ and a LOQ of 2.4 ± 0.16 ng mL⁻¹, the plant matrix produced a LOD of 4.5 ± 0.8 ng mL⁻¹ and an LOQ of 13 ± 2.6 ng mL⁻¹ (**Figure 5B**). This approximative 6-fold loss in sensitivity of the assay, while not negligible, remains compatible with robust quantitative assessment of dsRNA from crude plant extracts with Sand-BIRD.

We subsequently tested Sand-BIRD on healthy tissues and tissues infected with two positive-sense single-stranded RNA (+ssRNA) viruses. For this, we selected Tomato bushy stunt virus (TBSV) and Grapevine fanleaf virus (GFLV), which differ in their replication dynamics and capacity to produce dsRNA. A previous comparative study showed that TBSV replication leads to massive production of dsRNA, in contrast to GFLV, which produces amounts below the detection limit [9]. Analysis of crude extracts from TBSV-inoculated (TBSV-Inoc) and systemically infected *N. benthamiana* leaves (TBSV-Sys) revealed significantly elevated dsRNA levels compared to healthy controls (*p*-value <0.0001; n = 8) (**Figure 5C**). Parallel testing of these same samples with a commercial anti-TBSV ELISA (Bioreba) confirmed the infection status and the presence of TBSV in both inoculated and systemic leaves (**Figure 5C**). Moderately but significantly elevated dsRNA levels (*p*-value < 0.0001; n=8) were similarly detected in GFLV-infected plant extracts, with infection independently confirmed and quantified by anti-GFLV ELISA (Bioreba) (**Figure 5C**).

To further evaluate the Sand-BIRD platform’s capacity for disease monitoring, we conducted time-course studies tracking the progression of both TBSV and GFLV infection in plants over time. Following TBSV inoculation in *N. benthamiana*, the assay detected dsRNA in inoculated leaves from 3 days post infection (dpi) onward, reaching a plateau at 7 dpi. In systemic leaves, dsRNA was first detected at 6 dpi, peaked at 8 dpi, and subsequently declined to a stable plateau at 10 dpi (**Figure 5D**). This delayed dsRNA response in systemic leaves is consistent with the time required for long-distance vascular transport of the virus. This dsRNA response was compared with Bioreba anti-TBSV ELISA, which manifested a dose-dependent profile starting at 2 dpi and 4 dpi in inoculated and systemic leaves, respectively, and reached a plateau after 3 and 8 dpi (**Figure 5E**). For GFLV, dsRNA was first detected at 6 dpi in inoculated leaves and at 9 dpi in systemic leaves, coinciding with the appearance of symptoms in systemic leaves (**Figure S12A**). Across both pathosystems, the Bioreba virus ELISA signal consistently preceded dsRNA detection by approximately 2–4 days, reflecting the temporal offset between virion accumulation and peak dsRNA production. Collectively, these time-course experiments establish that Sand-BIRD faithfully captures viral replication dynamics through the sensitive detection of dsRNA, positioning it as a valuable tool for early and progressive monitoring of plant virus infections.

Having demonstrated Sand-BIRD’s ability to detect dsRNA in virus-infected plant matrices, we next assessed whether this compatibility extends to animal and human viral vectors. To this end, we performed a comparable spike-in validation using crude extracts from healthy *Aedes aegypti* mosquitoes as a control matrix (cHM). Under these conditions, Sand-BIRD achieved an LOD of 2.8 ± 0.2 ng mL⁻¹ and an LOQ of 8.5 ± 0.4 ng mL⁻¹ (**Figure 5F**). Together, these spike-in calibration curves demonstrate that the biosensor performs reliably in both plant and arthropod vector matrices without significant signal quenching, providing the necessary baseline for evaluating genuinely infected samples.

We next evaluated Sand-BIRD performance using flavivirus-infected mosquito samples (**Figure 5G**). Vector-borne flaviviruses represent a critical and expanding nexus of emerging infectious disease (EID) risk worldwide, making sensitive, field-deployable detection tools particularly valuable. Sand-BIRD successfully discriminated healthy mosquitoes from those infected with West Nile virus (WNV) or Dengue virus (DENV). When healthy and intrathoracically inoculated mosquito samples prepared in independent batches were tested, all uninfected control samples consistently fell below a baseline threshold of ∼0.2 corrected absorbance units. In contrast, independent biological replicates of WNV- and DENV-infected mosquitoes exhibited significantly elevated dsRNA levels (p = 0.0002–0.001 for WNV and p = 0.001–0.02 for DENV; n = 4), with the exception of a single DENV replicate that yielded a lower signal (**Figure 5G**). Active infection status in WNV- and DENV-infected samples was independently confirmed by RT-qPCR (**Figure S13**). Collectively, these results establish Sand-BIRD as a robust dsRNA-based surveillance platform capable of detecting arboviral infection across biologically distinct host matrices.

The epidemiological relevance of this capability is underscored by the escalating global burden of arboviral disease. Dengue alone accounted for approximately 14.1 million reported cases in 2024 - a twofold increase over 2023 and a 12-fold rise relative to 2014 - with roughly 9,000 associated deaths recorded worldwide [67]. WNV, though frequently asymptomatic, causes febrile illness in approximately 20% of infected individuals, and neuroinvasive complications, including meningitis and encephalitis, develop in approximately 1% of cases [68]. Together, these figures highlight an urgent and unmet need for rapid, sensitive, and deployable diagnostic tools capable of detecting arboviruses in mosquito vectors before human transmission occurs - a gap that Sand-BIRD is well positioned to address.

### Validation of Oxford Nanopore direct dsRNA sequencing

While Sand-BIRD dsRNA sensing provides critical quantitative evidence of viral presence, it does not allow direct identification of the originating pathogen. We therefore sought to develop a pipeline combining dsRNA sensing with nanopore direct dsRNA sequencing. As a first step, we validated the performance of Oxford Nanopore Technologies (ONT) direct RNA sequencing protocol (dRNA-seq) using two reference materials: (1) Larifan, a mixture of dsRNA from bacteriophage origin [69], and (2) a 142 bp dsRNA positive control of unknown sequence from Jena Bioscience (**Figure S14**). Larifan consists of dsRNA extracted from bacteriophage-infected *E. coli* cells, with heterogeneous lengths spanning approximately 50–5,000 bp [69], making it an excellent *bona fide* duplex RNA standard for the systematic calibration of dsRNA-specific workflows, library preparation, and bioinformatic analysis.

Since ONT does not natively support barcode multiplexing for direct RNA sequencing with the RNA004 chemistry, we adopted the synthetic barcode approach implemented in the WarpDemuX workflow [70]. WarpDemuX achieves signal-based demultiplexing through ligation of a unique identifier-containing sequence and represents one of the most accurate and efficient dRNA-seq barcoding strategies currently available, outperforming earlier tools such as DeePlexiCon [71] in terms of accuracy, speed, and computational resources. In this approach, the barcode sequence is embedded within a custom reverse transcription adaptor (RTA) [70]. Importantly, although the RTA is ligated to polyadenylated dsRNA for barcoding purposes, we omitted the reverse transcription step specified in the standard ONT dRNA-seq protocol. This decision is justified by the thermodynamic stability of the dsRNA duplex, which renders complementary strand synthesis prior to sequencing adaptor ligation unnecessary. Following poly-A tailing (**Figure S14A-B**) and adaptor ligation, ONT dRNA-seq was performed on 142 bp and Larifan dsRNA species on a single MinION flow cell (**Figure S14C**) using WarpDemuX barcodes 5 and 7 (WDX_RTA_BC5 and WDX_RTA_BC7, respectively), in what we refer to hereafter as the dsNanoSeq workflow.

Approximately 14% of high-quality reads (54,135 reads; WarpDemuX barcode prediction confidence (BpC) > 0.98; Phred quality score Q ≥ 10) were retrieved from the raw data pool for Larifan after filtering with Chopper (**Figure S15A**) and subsequently mapped to the bacteriophage MS2 genome using a combination of minimap2 and BLASTn (Basic Local Alignment Search Tool, for nucleotide). Of these, approximately 98% (53,057 out of 54,135 reads) mapped successfully to the MS2 genome, while only 29 reads mapped to the host *E. coli* genome (**Figure S15B**). Read length versus quality kernel density estimation (KDE) analysis revealed a population skewed toward shorter reads (N50 = 299 nt); however, the sequencing also captured longer dsRNA species exceeding 2 kb, with an average Phred quality of Q12 (**Figure S15C**). Reads assigned to the Larifan-specific barcode mapped across the full 3,569 nt MS2 genome (NC_001417.2), achieving a coverage depth of 2,000–6,000x, with balanced strand representation (forward: 22,508 reads, 44%; reverse: 28,480 reads, 56%) and complete genome coverage (**Figure S15D**).

For the 142 bp synthetic dsRNA control, read length versus quality KDE analysis generated by NanoPlot revealed a clear peak in the 120–160 bp region (**Figure S16A**). Alignment of reads within this cluster yielded an unambiguous consensus sequence matching the 142 bp positive control, which, as expected, is a fully synthetic RNA duplex species, and returned no significant similarity in a standard BLASTn search. Taken together, these results demonstrate that direct nanopore sequencing of dsRNA is achievable without a reverse transcription step, requiring only poly-A tailing prior to adaptor ligation and sequencing. The approach successfully captures full-length dsRNA species exceeding 3 kb at high coverage, with no prior knowledge of the dsRNA sequence required, establishing dsNanoSeq as a sequence-agnostic and RT-free framework for direct duplex RNA characterisation.

### Combining dsRNA detection with nanopore sequencing for virus identification

Having established the dsNanoSeq protocol on well-defined reference dsRNA standards, we next applied it to virus-infected plant and insect samples. The core concept underlying this integrated approach is to subject a Sand-BIRD-positive sample to a downstream nanopore sequencing pipeline by directly eluting captured dsRNA from the 96-well plate surface (**Figure 6A**). In this end-to-end workflow, the B2-mScarlet-coated surface serves a dual function: it acts simultaneously as a biosensing platform for dsRNA quantification and as an affinity enrichment matrix from which bound dsRNA can be eluted and channelled directly into sequencing for pathogen identification.

**Figure 6.**
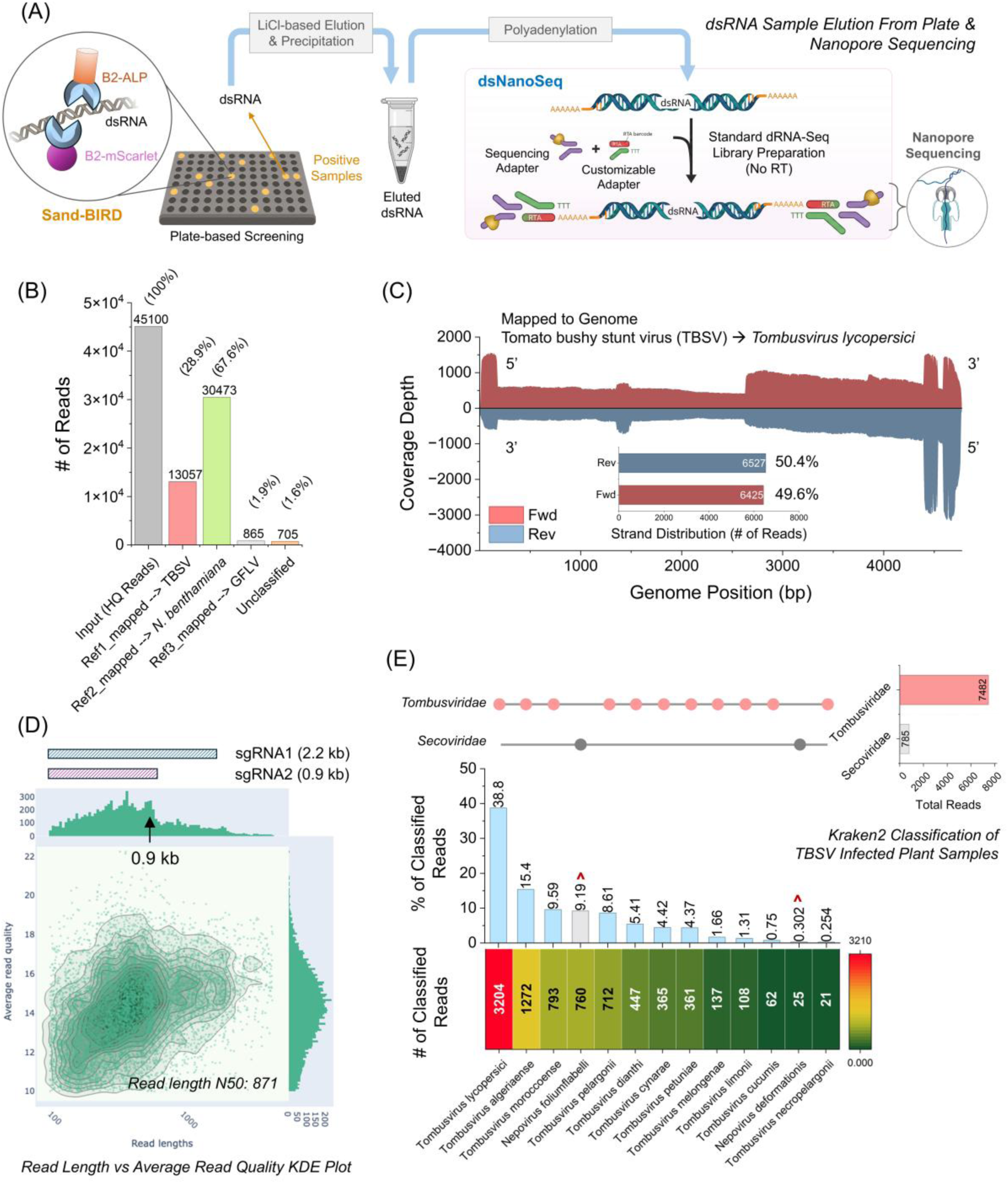
Nanopore direct dsRNA sequencing of the virus-infected samples. **(A)** Schematic illustration of the dsRNA elution from the 96-well microplate and nanopore sequencing. The samples appearing positive in terms of dsRNA level in Sand-BIRD were eluted from the B2-mScarlet surface, polyadenylated, and subjected to the Oxford Nanopore Technologies (ONT) direct RNA sequencing library preparation protocol using RNA004 chemistry. The dsRNA library was then processed for nanopore sequencing by either MinION or PromethION flow cells. **(B)** Mapping statistics of all the high-quality (HQ) reads from the TBSV-specific barcode. Around 29% of all the reads map to the TBSV genome, while 67% was classified as host *N. benthamiana* reads. **(C)** Genome coverage and profile of the reads mapping to the TBSV reference genome. The forward (red) and reverse (blue) strand reads cover the entirety of the genome in almost equal proportion (figure inset; 49.6% forward reads, and 50.4% reverse reads). **(D)** Read length vs average read quality distribution kernel density estimate (KDE) plot of the mapped TBSV reads as obtained from NanoPlot. The N50 of the read length is 871 nt, which is nearly similar to the TBSV subgenomic RNA2 (sgRNA2 peak in the distribution is indicated by an arrow). The size of the TBSV subgenomic RNA1 and RNA2 is displayed schematically at the top. **(E)** Kraken2 taxonomic classification of the HQ reads from the TBSV barcode in an agnostic manner. Nearly 90% of the reads map to the species belonging to only the *Tombusviridae* family, confirming the TBSV infection. The data is visualized by an UpSet-style plot. The absolute number of reads and percentage are detailed by the matrix plot and bar plot, respectively. The dots and connecting lines act as a map to the bottom x-axis to their broader viral families. Pink dots represent species within *Tombusviridae*, and grey dots represent species within *Secoviridae*. The top right bar plot displays the total absolute read counts assigned at the family level, revealing the dominance of *Tombusviridae* (7,482 reads). Besides, around 10% of the reads map to two different *Nepovirus* species (marked by red ‘^’).

We first validated dsRNA elution from Sand-BIRD wells loaded with TBSV-infected plant extracts and subjected the eluate to dsNanoSeq. We refer to this pipeline - combining Sand-BIRD-based dsRNA capture and quantification with downstream dsNanoSeq - as **BIRD-Seq** (B2-Integrated dsRNA Detection and Sequencing). For elution, wells containing TBSV-infected samples were washed and treated with high-salt buffer (2.5 M LiCl), after which dsRNA was coprecipitated with GlycoBlue carrier in isopropanol (see Methods). The recovered dsRNA was resuspended in nuclease-free water and subjected to poly-A tailing, and adaptor ligation before nanopore sequencing according to the dsNanoSeq workflow described above (**Figure S14C**). Following the extraction of high-quality reads (Q ≥ 10; BpC > 0.98) from the raw data pool, reads were mapped to the genome of the TBSV strain BS3Ng [72]. Mapping yielded robust viral read identification: 29% of all high-quality reads (13,057 reads) mapped to the viral genome, while 67% (30,473 reads) mapped to the host *N. benthamiana* genome (**Figure 6B**).

Viral reads covered the TBSV genome continuously and completely (100% genome coverage), with a strikingly balanced strand distribution (forward: 6,425 reads, 49.6%; reverse: 6,527 reads, 50.4%), consistent with the capture of dsRNA replication intermediates produced during active viral replication by the B2 bioreceptor (**Figure 6C, S17A, B**). The read length distribution further aligned with the expected size profile of TBSV subgenomic RNAs (sgRNAs), with a peak at approximately 0.9 kb corresponding to the anticipated size of TBSV sgRNA2. The population N50 of 871 nt (**Figure 6D**) is consistent with the previously reported preferential accumulation of sgRNA2 beyond 16 hours post-infection (hpi) in *N. benthamiana* [73], providing orthogonal biological validation of the sequencing output.

It was also observed that 865 reads (1.9% of the high-quality reads or 0.7% of the original extracted reads from the nanopore run) mapped to the GFLV-GHu genome (**Figure 6B**). This was likely due to the barcode misassignment during the demultiplexing, since we used the same flow cell for sequencing TBSV and GFLV samples (with WDX_RTA_BC3 and BC4 for TBSV, and WDX_RTA_BC5 and BC6 for GFLV). Similar observation was previously discussed in the WarpDemuX literature, wherein barcodes (WDX4-WDX6) achieved an accuracy of 99.5% with newer RNA004 chemistry, with the probable 0.5% misclassification rate [70]. We further looked into the HQ reads in an agnostic manner and subjected them to Kraken2 taxonomic classification against a locally indexed viral RefSeq nucleotide database for agnostic virus species identification. The vast majority of classified reads aligned to viruses of the *Tombusviridae* family (∼90% of all the Kraken2 classified reads), with TBSV (*Tombusvirus lycopersici*) as the overwhelmingly dominant species, accounting for 38.8% (3,204 reads) (**Figure 6E**). The sole exception was the minor fraction of reads (785 in total) attributed to members of the family *Secoviridae*, predominantly GFLV, originating from the same flow cell and likely reflecting to the barcode misassignment during demultiplexing (**Figure 6E**).

Intriguingly, further analysis of our nanopore sequencing data from TBSV-infected plant samples revealed an abundant population of defective interfering (DI) RNAs (**Figure S17C**). This was evident from a strongly non-uniform coverage profile exhibiting peaks at the 5’ terminus, within an internal region of the p92 open reading frame (ORF), and at the 3’ end, which was split up by two recurrent deletions (Δ157→1,345 and Δ1,464→4,394). The retained segments correspond closely to the four non-contiguous regions (I-IV) that define prototypical tombusvirus DI RNAs, in which region I derives from the 5’ UTR, region II resides within the polymerase gene, and regions III-IV map to the 3’-terminus [74,75]. Comparable representation of both genomic strands was observed, confirming that these are *bona fide* replicating molecules rather than degradation products [75,76]. The presence of deletion variants in our experiment (∼0.17–0.51 kb) is consistent with the earlier proposed stepwise-deletion model of DI RNAs biogenesis [75,76], further demonstrating that the B2 surface efficiently captures the entire replicating dsRNA population. Taken together, these findings indicate that, beyond virus identification through dsRNA detection, BIRD-Seq can serve as a discovery tool to investigate fundamental aspects of virus biology and replication.

Analysis of the GFLV-infected sample revealed that approximately 3% of the high-quality reads mapped to the GFLV-GHu genome (**Figure 7A**). The read length distribution showed two prominent peaks at approximately 7.3 and 3.8 kb, corresponding to the GFLV RNA1 and RNA2 genomic segments, respectively (**Figure 7B**). An N50 of 3,771 nt further confirmed that most of the captured reads span the full-length of RNA2, with high average read quality (Q14-Q18). The observed excess of RNA2 over RNA1 reads is consistent with earlier reports by Roy et al. [77], in which RNA2 abundance exceeded that of RNA1 by 10 to 100-fold. Mapping of viral reads yielded complete genome coverage for both segments, with a pronounced enrichment of forward-strand reads in each case (**Figure 7C**). This discrepancy likely reflects the intrinsically lower dsRNA accumulation of GFLV relative to the high-titre tombusvirus TBSV, which constrains the dsRNA pool available for B2-mediated capture.

**Figure 7.**
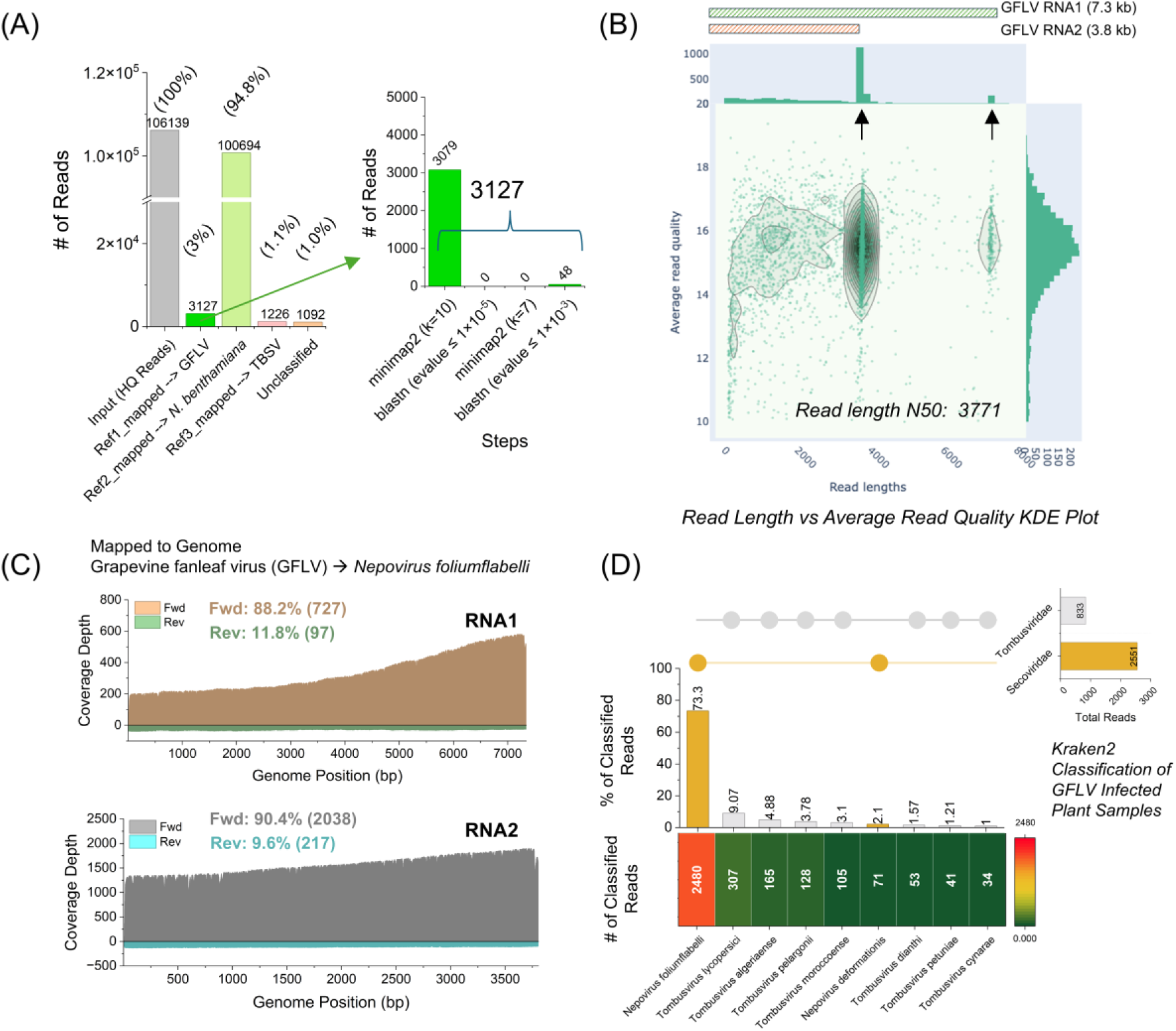
ONT direct dsRNA sequencing of GFLV-infected plant sample. (**A**) Statistics of the HQ reads mapping to the target GFLV GHu viral genome. Around 3% (3,127) of the reads were successfully classified as GFLV reads, validating the presence of the virus. (**B**) Read length vs. quality distribution (by NanoPlot) of the GFLV reads. Two clear peaks were identified around 3700-3800 nt and 7,200-7,300 nt (indicated by arrow), which represent the GFLV bipartite RNA genome (RNA1 ∼7.3 kb and RNA2 ∼3.8 kb; schematically shown at the top of the plot). (**C**) Genome coverage map of the viral reads of both RNA1 and RNA2. Around 100% of breadth coverage is achieved with a strong bias for forward reads (88% for RNA1 and 90% for RNA2). (**D**) Taxonomic classification of all the HQ reads in an agnostic manner (by Kraken2). The data is represented in an UpSet-style plot as described previously. Most of the reads (75%) are classified into the GFLV virus family *Secoviridae* (yellow bars), as expected. Around ∼25% reads were classified to *Tombusviridae* because of the barcode demultiplexing misclassification (TBSV and GFLV samples were sequenced on the same flow cell).

Agnostic taxonomic classification using Kraken2 assigned approximately 74% of reads to GFLV (*Nepovirus foliumflabelli*) (**Figure 7D**). As observed for TBSV, a minor proportion of reads were attributed to other members of the family *Tombusviridae*, consistent with barcode misassignment during demultiplexing, as discussed above. Taken together, BIRD-Seq identified members of the families *Tombusviridae* and *Secoviridae* exclusively in these samples, as expected given that the plant material had been inoculated with TBSV or GFLV, respectively.

To further validate the applicability of the BIRD-Seq, we analysed the WNV-infected mosquito samples, which inherently produced substantially lower dsRNA detection signals (in terms of absorbance) compared to TBSV or GFLV-infected plant samples (**Figure 5E**). Despite this, 3838 reads mapped successfully to the WNV genome, with the read length distribution centred in the 100-300 nt range (**Figure 8A-B**). Flavivirus RNA synthesis is inherently asymmetric, strongly favouring positive-sense genomic RNA over negative-sense replicative intermediates [78], which likely accounts for the pronounced strand bias observed in our data - despite achieving achieved 100 % genome coverage, forward-strand reads dominated the mapped read population (**Figure 8C**). The enrichment of shorter reads and pronounced accumulation of mapped reads at the 3’ end of the genome most likely reflect the abundant production of subgenomic flaviviral RNAs (sfRNAs) - nuclease-resistant RNA species generated by incomplete Xrn1-mediated 5’→3’ degradation of the WNV genome and known to accumulate to high levels during flavivirus infection [106]. The overall lower dsRNA yield in mosquito samples relative to plant hosts may additionally reflect differences in host antiviral RNAi activity, though this remains to be formally investigated. Agnostic taxonomic classification using Kraken2 identified a single viral species from the WNV-specific barcode (**Figure 8D**): all successfully classified reads were assigned to WNV (*Orthoflavivirus nilense,* 3,466 reads), unambiguously confirming the identity of the infecting virus. Collectively, these results demonstrate the capacity of BIRD-Seq to detect and identify arboviruses in mosquito vectors even when viral dsRNA levels are substantially lower than those encountered in plant hosts [79].

**Figure 8.**
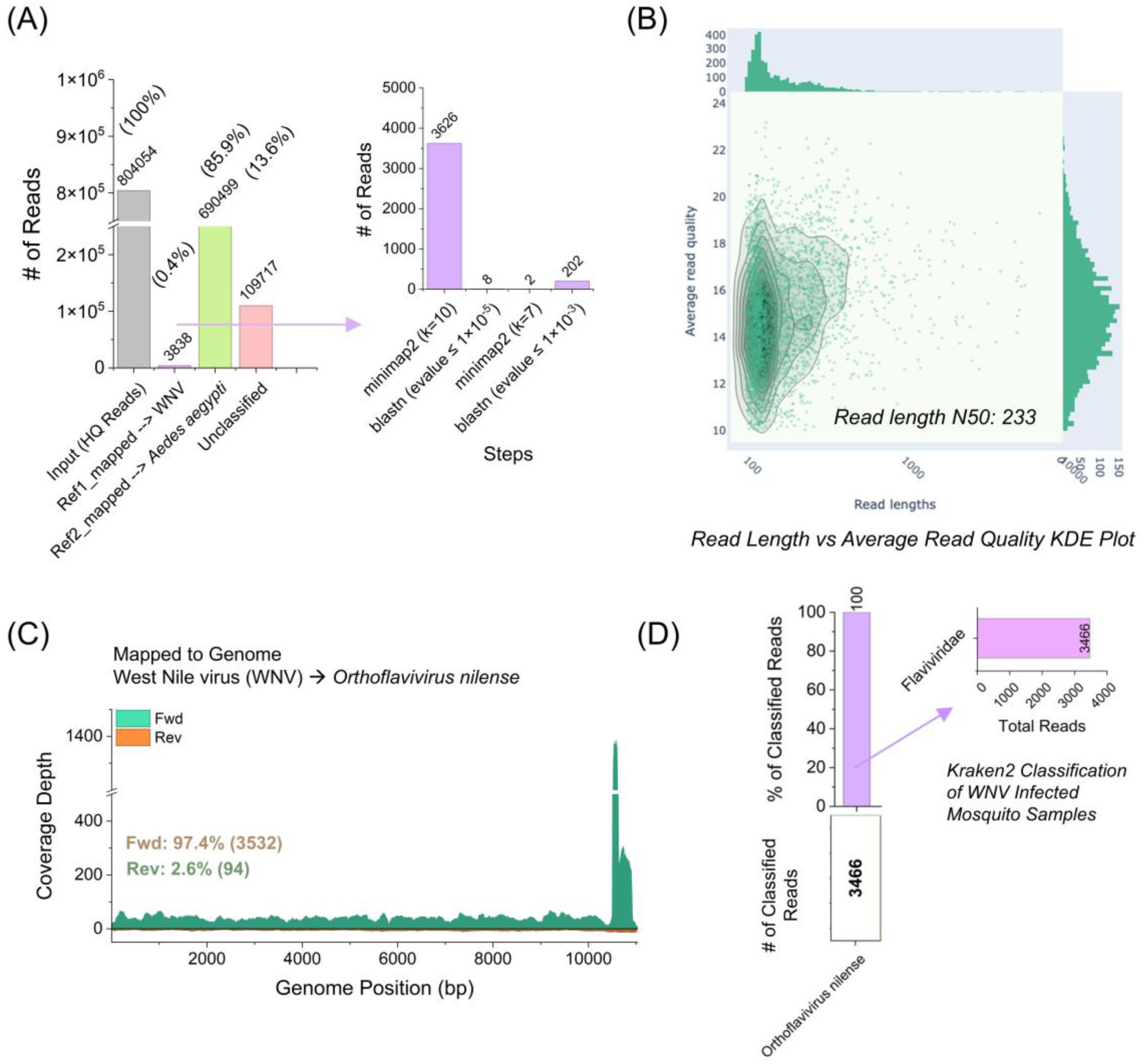
Nanopore direct dsRNA sequencing of WNV-infected mosquito sample. (**A**) Viral reads mapping statistics showcase around 3,838 reads map to the West Nile virus genome, ascertaining the infected virus species. (**B**) The read length distribution of the WNV reads as obtained from NanoPlot. The N50=233 suggests that the sequencing yielded much shorter reads in comparison to TBSV or GFLV. (**C**) Genome coverage mapping of the WNV reveals a high bias for forward-strand reads. Although a 100% genome coverage is achieved from the mapping of the sequencing reads, the 3’ end of the genome manifested much higher coverage depth. (**D**) Kraken2 taxonomic classification successfully identified (3,466 reads) only one virus species from the entire HQ reads population, mapping to the species *Orthoflavivirus nilense* (West Nile virus).

## Discussion

A sensitive, robust, and readily scalable dsRNA detection method holds considerable value not only for diagnostics and therapeutics, but also for fundamental research, given its broad applicability across the vaccine industry, virus diagnostics, and the study of innate immunity. Here, we developed a simple, robust, and mAb-free BIRD-Seq pipeline that couples extraction-free, quantitative dsRNA detection in complex matrices with Oxford Nanopore Technology (ONT) direct RNA sequencing for agnostic identification of viruses from both plant and mosquito samples. The BIRD-Seq pipeline directly addresses the principal limitations of mAb-based approaches by leveraging the B2 dsRNA-binding protein from Flock House virus (FHV) as a versatile bioreceptor, bridging quantitative dsRNA sensing with unbiased downstream pathogen identification via direct RNA sequencing.

To accurately profile B2/dsRNA binding kinetics, we developed the Res-BIRD surface plasmon resonance (SPR) platform. In this setup, B2 is first anchored to the sensor surface to capture long, natural-origin viral dsRNA. The multivalent, avidity-driven interaction between these long dsRNA molecules and multiple immobilized B2 sites creates a ‘velcro-like’ effect that arrests dissociation, yielding a highly stable, saturated RNA duplex layer. Using this uniform dsRNA surface, we characterized the binding kinetics of dsRNA-binding bioreceptors — including free B2 protein and the J2 mAb — by solution-phase titration. The equilibrium dissociation constant (*K*_D_) of B2 for *bona fide* viral dsRNA was determined to be in the low nanomolar range (8.6 nM), directly comparable to the gold-standard J2 mAb (*K*_D_ = 6.4 nM) measured under identical conditions. This affinity is also consistent with the recently described M2/M5 antibody pair, which exhibited *K*_D_ values of 9.45 nM and 5.38 nM, respectively, against a 40 bp dsRNA target [80]. While an earlier study reported an EMSA-derived *K*_D_ of 1.4 nM for an MBP-B2 fusion protein interacting with short dsRNA species [39], the Res-BIRD platform provides genuine kinetic resolution of the B2 dsRNA-binding domain engaging long, heterogeneous viral dsRNA (50–5,000 bp), which more faithfully recapitulates the biologically relevant substrate.

A major structural advantage of the B2 bioreceptor over conventional mAbs is its compact, all-alpha-helical, 17 kDa head-to-tail homodimeric architecture, which enables efficient and cost-effective production in bacterial expression systems. Moreover, its structural plasticity is amenable to modular bioengineering: B2 was seamlessly fused to heterologous reporter enzymes — mScarlet, NanoLuc luciferase, and alkaline phosphatase — without significantly compromising dsRNA-binding activity. The resulting Dot-BIRD assay operates as a streamlined, single-reagent detection equivalent to traditional two-antibody J2-based dot blots [81], achieving a comparable sensitivity of 0.2 ng per dot for viral dsRNA. Importantly, B2 fusion proteins exhibited a marked binding preference for natural viral dsRNA over synthetic structural analogues such as Poly(I:C) and Poly(A:U), and were practically non-responsive to ssRNA or dsDNA. This stringent selectivity validates the engineered B2 fusion proteins as reliable bioreceptors for natural viral dsRNA.

Transitioning to a quantitative, high-throughput format, the solid-phase sandwich dsRNA detection platform (Sand-BIRD) achieved a LOD of approximately 0.7 ng mL^-1^ in both the B2-ALP and B2-NanoLuc configurations. This performance is highly competitive with currently available commercial immunoassays, including the KRIBIOLISA K1-based kit (assay range: 10–640 ng/mL) and the GenScript J2-based ELISA (linear range: 0.123–30 ng mL^-1^). Critically, Sand-BIRD is functional in complex biological matrices without prior nucleic acid extraction — a capability absent from enrichment-dependent methods such as cellulose chromatography and DRB4-based kits or anti-dsRNA mAb-based approaches [11], which require purified total RNA as input. Sand-BIRD yielded low nanogram-per-milliliter sensitivities directly in unpurified, polyphenol-rich plant sap and protein-rich mosquito lysates, enabling the successful tracking of Tomato bushy stunt virus (TBSV) and Grapevine fanleaf virus (GFLV) infection kinetics from crude extracts. Notably, while earlier B2-based northwestern blotting failed to detect low-titer GFLV [9], Sand-BIRD successfully detected the virus in crude systemic leaf extracts as early as six days post-infection. The practical significance of this is underscored by the economic consequences of plant viruses. Globally, viral infections in plants are estimated to cause $30 billion in annual economic losses [83] [84], and the majority of certified diagnostic techniques remain virus-specific, relying on ELISA or RT-PCR. By contrast, the B2-based sandwich assay offers a generic, rapid, and sequence-agnostic front-end platform for broad-spectrum plant virus surveillance. The assay also reliably differentiated healthy mosquitoes from those infected with West Nile virus (WNV) and Dengue virus (DENV) at a 1:10 (w/v) tissue dilution, demonstrating its efficacy for scalable vector-borne disease monitoring.

Beyond its diagnostic utility, the Sand-BIRD platform features a unique dual-function modularity: it acts simultaneously as a physical enrichment matrix for downstream viral discovery. Standard metatranscriptomic approaches, while powerful, are frequently overwhelmed by host ribosomal and messenger RNA backgrounds, necessitating intensive total RNA extraction and costly host-specific ribodepletion [85]. In BIRD-Seq, selective capture of replicating viral dsRNA on the B2-mScarlet assay surface passively filters host background. For the positive samples, the captured RNA duplexes are then eluted directly and subjected to ONT direct RNA sequencing, substantially reducing host interference. For the high-titer model virus TBSV, this streamlined workflow recovered viral reads comprising approximately 29% of the high-quality reads, achieving considerable viral enrichment while entirely bypassing total RNA extraction and ribodepletion.

To contextualize the BIRD-Seq pipeline within the broader landscape of dsRNA-targeted surveillance, it is instructive to compare it with recent methodological advances. The strategy of coupling dsRNA enrichment with high-throughput sequencing has precedent in approaches such as dsRNA-Seq [12], which employs J2 mAb immunoprecipitation prior to Illumina sequencing, and NanoViromics [13], which demonstrated the value of combining dsRNA enrichment with long-read nanopore sequencing for agnostic plant virus detection. While these methods effectively enrich for actively replicating viral genomes and reduce host background, they depend on prior extraction of total RNA and multi-step protocols that frequently involve solvent-based extractions, single-strand-specific nuclease treatments, and ribodepletion. On the diagnostic side, solid-phase platforms employing reactive polymer surfaces immobilized with J2 antibody have demonstrated sensitive dsRNA detection from cell lysates [86], but rely on costly monoclonal antibodies and lack integrated downstream sequencing capacity. A PKR-based luminescent biosensor has also been reported to quantify viral dsRNA directly from crude cell culture lysates following detergent-based phase separation [45]; however, this approach does not provide an agnostic virus identification step. By substituting the J2 mAb with the modular, recombinant B2 protein and directly coupling an extraction-free capture assay to nanopore sequencing, BIRD-Seq eliminates the RNA purification and enzymatic digestion requirements of current enrichment methods, converting a crude-sample diagnostic into a comprehensive platform for pathogen discovery. To our knowledge, BIRD-Seq is the first integrated pipeline to simultaneously detect and quantify dsRNA directly in crude biological samples while providing sequence-independent virus identification through direct nanopore RNA sequencing. The complete BIRD-Seq workflow — from dsRNA sensing to sequencing — can be accomplished within 30 hours (<2 days; **Figure 9**), while the Sand-BIRD detection step alone requires only around 180 to 210 minutes from crude sample to result (excluding the coating step). Furthermore, the use of the SQK-RNA004 direct RNA sequencing chemistry means that BIRD-Seq analyses viral dsRNA strands as they exist in infected cells: positive- and negative-sense viral RNAs are treated equivalently, no reverse transcription step is required (eliminating a major source of sequence errors), and RNA base modifications (epitranscriptomic marks) are retained on viral strands as they traverse the nanopore, making them available for detection via the Dorado basecaller.

**Figure 9.**
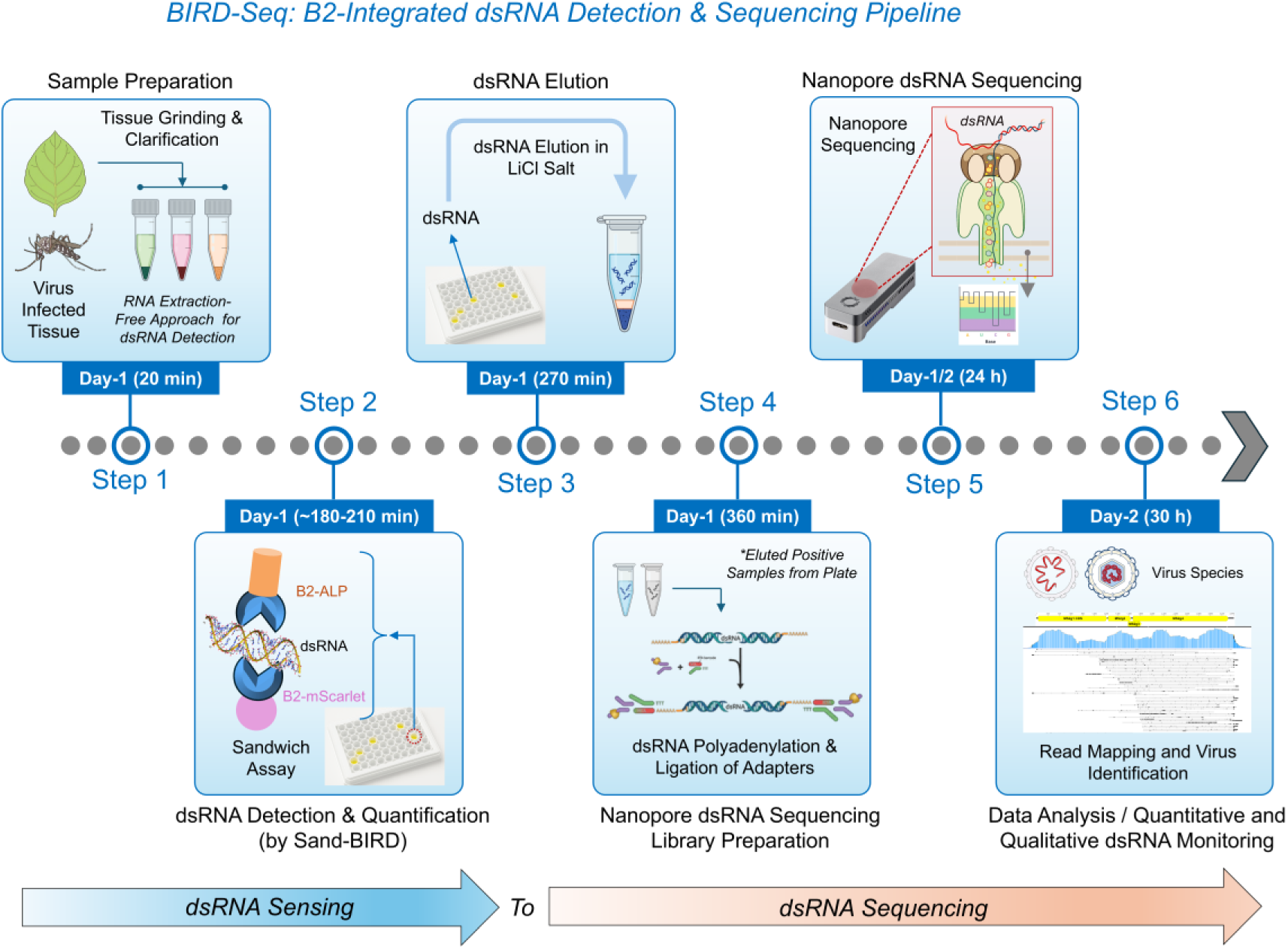
Schematic representation of the B2-Integrated dsRNA Detection & Sequencing Pipeline. The B2-implemented sandwich assay (Sand-BIRD) can be used for dsRNA-centred virus detection in crude samples within 3 hours. Furthermore, the whole end-to-end dsRNA sample-to-sequencing pipeline can be achieved in <2 days (∼30 h). Approximate time point necessary for each step is indicated within the schematic.

The BIRD-Seq pipeline, as currently implemented, has important limitations that merit careful consideration. Sand-BIRD robustly captures active viral replication, but it cannot resolve individual viral species in mixed infections without the downstream dsNanoSeq step; it should therefore be viewed as a highly sensitive, sequence-agnostic screening platform rather than a standalone diagnostic. Although the assay performs comparably to commercial ELISAs in crude extracts, very early or low-titer infections may still require confirmatory target-specific RT-qPCR. A signal below the established baseline should thus be interpreted as dsRNA remaining below the assay’s detection threshold, not as definitive evidence against active infection. Deployment in on-site diagnostic settings will also require a rigorous, matrix-specific baseline established from healthy crude extracts before screening, because viral replication kinetics and dsRNA accumulation vary substantially across viral species, tissue types, and infection stages. Importantly, B2 should not be regarded as the ultimate non-mAb-based dsRNA sensor, given the wider diversity of host- and virus-encoded dsRNA-binding proteins with distinct affinities, topologies, and biological roles. While FHV B2 proved highly effective in the present study, the structural diversity of natural dsRNA-binding proteins represents a rich space for further exploration. Alternative bioreceptors, including viral suppressors of RNA silencing such as NS1 and VP35, or host innate immune sensors such as PKR and the plant DRB protein family, may yield additional high-affinity, high-selectivity dsRNA-binding scaffolds. Finally, although Sand-BIRD offers a cost-effective screening step, the nanopore direct RNA sequencing stage remains relatively expensive and requires sufficient dsRNA input; moreover, the platform is intrinsically restricted to actively replicating viruses because it detects replication intermediates and will not identify latent or dormant pathogens.

Future developments of the platform could adapt the B2 bioreceptor to on-site rapid testing formats, such as lateral flow assays, and evaluate its performance in clinical mammalian matrices such as serum or saliva. Notably, circulating dsRNA has been detected at nanogram-per-milliliter levels in the serum and plasma of patients with viral infections and inflammatory diseases such as rheumatoid arthritis and multiple sclerosis [107], while the presence of extracellular dsRNA in saliva remains essentially unexplored, underscoring the potential diagnostic value of deploying a sensitive, matrix-tolerant platform such as Sand-BIRD in these clinical fluids.

## Concluding Remarks

The BIRD-Seq platform demonstrated in this work aligns with the World Health Organization’s One Health framework, which recognizes the interconnection between plant, animal, and human health and calls for integrated, cross-kingdom surveillance strategies. Double-stranded RNA, as the universal molecular hallmark of RNA virus replication, represents an ideal agnostic target for such broad-spectrum surveillance. The B2-protein–based Sand-BIRD assay is highly robust in complex crude matrices derived from both plants and insects, achieving nanogram-per-milliliter sensitivity without prior nucleic acid extraction, and constitute a compelling alternative to J2-based immunodetection at substantially reduced production cost. Proof-of-concept virus surveillance was rigorously demonstrated across a phylogenetically diverse panel of RNA viruses spanning two kingdoms: the plant viruses, such as, Tomato bushy stunt virus and Grapevine fanleaf virus, and the arboviruses like West Nile virus and Dengue virus.

Beyond detection, we present a simple and efficient protocol for the direct elution of captured dsRNA from the B2-coated assay surface, enabling seamless integration of the Sand-BIRD quantification step with direct dsRNA sequencing. This streamlined BIRD-seq workflow allows dsRNA from positive samples to be eluted from the assay plate and directly subjected to Oxford Nanopore sequencing without prior RNA extraction steps, and was validated for TBSV, GFLV, and WNV, yielding strand-resolved, quantitative information on the viral dsRNA species present. To our knowledge, BIRD-Seq represents a fully integrated pipeline in which multiple crude biological samples can be processed in parallel, simultaneously quantifying dsRNA by sandwich immunoassay and identifying the causative viral agent by direct nanopore sequencing and thereby offering a scalable, cost-effective, and sequence-agnostic solution for emerging infectious disease surveillance. Future iterations will focus on improving viral read recovery in low-titer scenarios and on large-scale validation using field-collected samples from diverse host species and geographic contexts.

## Materials and Methods

### Materials

The tris(2-carboxyethyl)phosphine) (TCEP, Cat. 580560), PEG (Poly(ethylene glycol) methyl ether thiol, Cat. 729108), Phosphate-buffered saline (PBS) tablets, Tween-20, NaCl, HEPES, and antibiotics were purchased from Sigma-Aldrich. The viral dsRNA (Larifan; referred to as V-dsRNA in this article) was kindly provided by Dr. Dace Pjanova, Riga Stradins University, Latvia. All the V-dsRNA samples were prepared from a stock of 30 µg mL^-1^ (in 1× PBS) and quantified by the Thermo Fisher Scientific NanoDrop instrument. All the enzymes required for cloning and PCR were purchased from New England Biolabs (NEB). The DNA oligonucleotides (PCR primers) and gene blocks were obtained from Integrated DNA Technologies (IDT). The J2 antibody and 142 bp standard control dsRNA were purchased from Jena Bioscience. The Poly(I:C), Poly(A:U), and ssRNA (Poly(U)) were purchased from InvivoGen. The pNPP substrate (p-Nitrophenyl Phosphate) for B2-ALP is purchased from J & K SCIENTIFIC LTD (Cat. 254303). The furimazine substrate was purchased from CliniSciences. This product has since been discontinued; furimazine is available from Promega (e.g., Nano-Glo luciferase assay substrate). All the terminologies – ‘V-dsRNA’, ‘Larifan’, ‘*bona fide* viral dsRNA’, ‘dsRNA of natural origin’-are used interchangeably in this article.

### Software

The modelling of all the proteins, wherever applicable, was done by the AlphaFold-3 webserver (https://alphafoldserver.com/) [90]. The structures were then downloaded and visualized either by PyMOL (Version 3.0, Schrödinger, LLC) or ChimeraX (v1.11.1, UCSF). In most cases, multiple models with various seed numbers (seed 1-5) were generated to substantiate the validity of the models. A list of all the protein constructs is provided in **Table S1**. All the primer design and in silico cloning experiments were done by Geneious (https://www.geneious.com/). All the graphs/plots presented in this article were produced by either OriginLab (https://www.originlab.com/), GraphPad (https://www.graphpad.com/), or custom-written R scripts. The illustrations were either taken from open-source NIAID NIH BioArt (https://bioart.niaid.nih.gov/), prepared by Affinity Designer, or MS PowerPoint.

### Cloning

B2-SpT and C-SpyC were cloned into a plasmid suitable for heterologous expression in the cytoplasm of *Escherichia coli* by Golden Gate assembly into the pET-GG (cyto) vector, while B2-ALP was cloned into the pET-GG (peri) vector for periplasmic expression. Both pET-GG vectors are pET-22 derivatives adapted for Golden Gate cloning using the type IIS restriction enzyme SapI. The B2 module was generated by PCR amplification of the sequence encoding the first 72 amino acids of B2 (GenBank accession X77156), flanked by SapI-compatible sites. The ALP module was amplified from plasmid pLip6v, which contains the genomic sequence of bacterial alkaline phosphatase, using primers that introduced a C-terminal 6×His tag and SapI-compatible sites. The pLip6v was a kind gift from Bruno Müller [62,63]. The SpT module was generated from a SpyTag003-containing template using primers that introduced a 3×(SGGG) linker between B2 and SpyTag003, a C-terminal 6×His tag, and SapI-compatible sites. B2-SpT was obtained by Golden Gate assembly of the B2 and SpT modules into pET-GG (cyto), whereas B2-ALP was obtained by assembling the B2 and ALP modules into pET-GG (peri). Cys-SpyCatcher003 (C-SpC) was cloned into the pET-GG (cyto) vector by the assembly of the 6His-Cys-3GS3G module and the SpyCatcher (SpC) PCR module. The 6His-Cys-3GS3G module was ordered as a synthetic gene from Integrated DNA Technologies and contains a 6×His tag, a single cysteine residue, a 3GS3G linker, and SapI-compatible sites. The SpC module was generated using primers that introduced SapI-compatible sites.

B2-NLuc and B2-mSc were cloned into a pDGB_Ω-derived expression vector by GoldenBraid assembly of a B2 module with NLuc and mScarlet modules, respectively. The B2 module was obtained as described above, except that BsaI-compatible sites were used. NLuc, containing an additional C-terminal 6×His tag and BsaI sites, was obtained as a synthetic gene from Integrated DNA Technologies. The mScarlet sequence, with a C-terminal 6×His tag, was codon-optimized for E. coli expression and ordered as a synthetic gene carrying BsmBI restriction sites, then cloned into the Level 0 pUPD2 vector. The mScarlet tag (mScarlet-I, FPbase 6VVTK) was chosen because it lacks cysteine residues, which is advantageous for fusion to cysteine-containing proteins by limiting unwanted disulfide bond formation, misfolding and aggregation. Assemblies of the different modules were performed using a 3:1 molar ratio of agarose gel-purified PCR products or synthetic gene to 150 ng of pET-GG (cyto) plasmid in a reaction containing 10 units of SapI or BsaI-HFv2 (New England Biolabs), 1× rCutSmart buffer (New England Biolabs), 5 units of T4 DNA ligase (Thermo Scientific), and 0.5 mM ATP (Thermo Scientific). The reaction was subjected to 30 cycles of 37 °C for 10 min and 18 °C for 10 min, followed by incubation at 18 °C for 1 h, 50 °C for 10 min, and 80 °C for 10 min. Plasmids were transformed into *E. coli* TOP10 chemically competent cells by heat shock and selected on carbenicillin (100 µg mL⁻¹). Colonies were grown in liquid LB medium containing carbenicillin, plasmids were purified using the NucleoSpin Plasmid QuickPure kit (Macherey-Nagel), and the integrity of all inserts was confirmed by Sanger sequencing.

### Recombinant Protein Expression and Purification

Recombinant B2 fusion proteins (B2-nLuc, B2-ALP, and B2-mScarlet) were expressed in *E. coli* BL21 (DE3) using auto-induction medium [91]. Briefly, 50–100 ng of plasmid DNA was transformed into thermocompetent BL21 cells by heat shock, recovered in LB medium, and used to inoculate 50 mL LB starter cultures supplemented with 2% glucose and 100 µg mL⁻¹ carbenicillin. After overnight growth at 37°C, starter cultures were used to inoculate 900 mL of auto-induction medium (NZY base supplemented with 20× NPS salts, 50× 5052 solution, MgSO₄, and trace elements) in 2.5 L Ultra Yield flasks (Generon) to a starting OD₆₀₀ of 0.05. Cultures were incubated at 37°C for 4 h and then shifted to 20°C overnight to induce protein expression. Cells were harvested by centrifugation (4,000 × g, 8 min, 4°C), resuspended in lysis buffer (50 mM NaCl, 50 mM HEPES, pH 7.5, 1× protease inhibitor cocktail), and disrupted by microfluidization (2 passages, 1,305 bar). The lysate was clarified by centrifugation (17,000 × g, 2 h, 4°C) and filtration (0.22 µm). The supernatant was supplemented with 25 mM imidazole and loaded onto a HisTrap™ FF 5 mL column (Cytiva) pre-equilibrated in binding buffer (50 mM HEPES, pH 7.5, 1 M NaCl, 25 mM imidazole). Bound proteins were eluted with imidazole gradient up to 500 mM and further purified by size exclusion chromatography on a HiLoad™ Superdex™ 200 16/600 column (Cytiva) equilibrated in 50 mM HEPES, pH 7.5, 300 mM NaCl. Fractions containing B2 fusion proteins were pooled, analyzed by SDS-PAGE, and stored at −80°C.

### Surface Functionalization and B2 Immobilization for SPR

A gold-coated SPR prism (obtained from Affinité Instruments, Canada) was first coated with the Cys-SpyCatcher003 (C-SpC) protein. For this, 40 µM of C-SpC (in 1× PBS-T buffer: 1× PBS, 0.05% v/v Tween-20) was mixed with 20 mM of TCEP in a 1:1 v/v ratio, incubated at room temperature (RT) for 5 min, and drop-casted over the gold prism. This mixture was then incubated at RT for 2 hours, which allowed the immobilization of the C-SpC to the gold surface by the thiol (-SH) linkage. The prism was subsequently rinsed gently with 1× PBS-T and MiliQ water. Next, the prism was PEGylated to incorporate antifouling properties. Poly(ethylene glycol) methyl ether thiol (10 mM) (Sigma, Cat. 729140) was mixed with TCEP (20 mM) in a 1:1 v/v ratio, drop-casted over the prism, and incubated for 30 min at RT. The prism was then rinsed gently with 1× PBS-T and MiliQ water again before inserting it into the SPR instrument. The B2-SpyTag003 (B2-SpT) protein immobilization (over the C-SpC surface) was further carried out inside the SPR instrument. 2 µM of B2-SpT was injected for 6 mins, incubated on the prism for 5 mins, and then washing was carried out (**Figure S2B**). For the dsRNA titration and sensing experiment, a series of different concentrations of V-dsRNA (5-3,000 ng mL^-1^) was injected on the B2 functionalized prism. Next, the kinetics of the B2 and all engineered versions (B2-mSc, B2-ALP, B2-NLuc) were retrieved by injecting them (50, 100 nM) over a saturated viral dsRNA (V-dsRNA) surface (created by flowing 500 ng mL^-1^ V-dsRNA over the B2-coated interface). The association and dissociation of all protein constructs are recorded in this way, reflecting their binding affinity towards a dsRNA.

### Surface Plasmon Resonance (SPR) Measurements

SPR measurements were performed on a portable P4 SPR platform from Affinité Instruments, Canada. The resolution of the detector was 1.34 nm full width at half maximum (FWHM). The P4SPR contains 4 parallel channels (referred to as A-D) for analyzing the samples, and for each kinetics study, one of the channels was kept as ‘reference’ (only buffer flow). SPR curves were recorded using a running buffer of 1× PBS-T (1× PBS, 0.05% v/v Tween-20) at a flow rate of 30 µL min^-1^. Surface modification of the gold-coated prims was performed as per the protocol mentioned earlier. All binding kinetics experiments in this study were conducted at room temperature, and all sensorgrams were fitted using GraphPad Prism software – ‘association and dissociation kinetics’ model (Version 8, GraphPad Software, Boston, Massachusetts, USA, https://www.graphpad.com).

The limit of detection (LOD) and limit of quantification (LOQ) parameters throughout the work were determined by using: LOD = (3 × B_SD_) / S and LOQ = (10 × B_SD_) / S, respectively, wherein the B_SD_ = standard deviation of the blank, S = slope of the lower linear region of the calibration curve [92]. Statistical significance between the two experimental groups, wherever applicable, was calculated using Welch’s two-sample t-test (assuming unequal variances) performed in Microsoft Excel.

### B2-based Dot Blot for dsRNA Monitoring

Dot blot assays were adapted from standard nucleic acid detection protocols using B2 fusion proteins to assess the binding of different nucleic acid species. Hybond-N⁺ membranes (GE Healthcare) were spotted with serial fourfold dilutions of nucleic acids (0.05 - 200 ng µL⁻¹; 20 µL per dilution) prepared in 96-well PCR plates. Six nucleic acid types were tested across seven rows, with the eighth row reserved as a negative control (no nucleic acid or no template). Aliquots of 2 µL from each dilution were applied in triplicate to three replicate membranes, air-dried, and UV cross-linked prior to downstream processing. **Figure S9** provides a schematic pipeline of the method.

1. B2-ALP Dot Blot: Membranes were blocked for 2 h in 5% milk in PBS containing 0.1% Tween-20, then incubated overnight at 4°C with B2-ALP (0.25 µg mL⁻¹ in blocking buffer). After three washes in PBS-T (5 min each), membranes were equilibrated for 2 min in conjugate buffer (2.4 g Tris, 8 g NaCl, 20 g PVP K25, 0.5 g Tween-20, 2 g BSA, 0.2 g MgCl₂·6H₂O, 0.2 g KCl, 0.2 g NaN₃ per liter, pH 7.4) and developed with BCIP/NBT liquid substrate (Thermo Scientific) for ∼10 min in the dark. Reactions were stopped by rinsing in water, and membranes were photographed for dot visualization.
2. B2-NLuc Dot Blot: Membranes were blocked for 2 h in 5% milk in PBS containing 0.1% Tween-20, then incubated overnight at 4°C with B2-nLuc (0.5 µg mL⁻¹ in blocking buffer). After four washes in PBS-T (5 min each), membranes were briefly equilibrated in substrate buffer (100 mM MES, 1 mM EDTA, 150 mM KCl, 35 mM thiourea, pH 6) and exposed to 10 µM furimazine for ∼4 min in sealed plastic sleeves. Luminescence was recorded on a Fusion FX imaging system (Vilber), and signal intensities were quantified using ImageJ (NIH) Gel Analysis. Concentration vs. response relationships were plotted, and all experiments were repeated in triplicate to ensure reproducibility.

### B2-based Sandwich Assay for dsRNA Detection (Sand-BIRD)

#### Principle of Sand-BIRD

The core idea and working of the B2-integrated sandwich assay (Sand-BIRD) was essentially inspired by the conventional double antibody sandwich ELISA (DAS-ELISA), wherein two sets of antibodies specific for an analyte are used for the detection purpose. One set of antibodies is employed for coating the plate (essentially facilitating the analyte capturing), and another set coupled to a reporter enzyme is used for revealing the bound molecules. For Sand-BIRD, B2-mScarlet (B2-mSc) was used for coating, and this surface captures the dsRNA species. Then either B2 alkaline phosphatase (B2-ALP) or B2 fused to NanoLuc luciferase (B2-NLuc) was used to reveal the presence of dsRNA. The pipeline of the assay is illustrated in **Figure 4A**. The assay is validated for both absorbance (B2-ALP, p-Nitrophenyl Phosphate (pNPP) as substrate) and a luminescence (by using B2-NLuc, furimazine as a substrate)-based approach.

#### Protocol for the Sand-BIRD

Three different types of 96-well MaxiSorp plates from Thermo Fisher Scientific were used for the sandwich assay. Clear transparent plates (Cat. 442404) were used for the B2-ALP; black (Cat. 437111) or white (Cat. 436110) plates were used for the B2-NLuc. The plates were first coated with B2-mScarlet (7 µg mL^-1^) prepared in coating buffer (composition for 1,000 mL, pH 9.6: Na₂CO₃ 1.59 g, NaHCO_3_ 2.93 g; add 1 mM DTT right before the B2-mSc preparation) for overnight at 4°C. Plates were then washed with wash buffer (WB: 1× PBS, 0.1% Tween-20) for 3 times. The blocking of the plates was done by adding 300 µL blocking buffer (BB: 1× PBS, 5% w/v non-fat dry milk (NFDM), 0.1% v/v Tween-20) to each well, and the plates were incubated at 28°C for 1 h. Washing of the plates 3 times by WB was done afterwards, before adding samples to be analysed for dsRNA. For performing the standard calibration curves, a *bona fide* viral dsRNA (V-dsRNA) sample was used, which was a heterogeneous dsRNA population of 50-5,000 bp (mean molecular mass of about 500 kDa). The V-dsRNA sample was isolated from the bacteriophage MS2-infected *E. coli* cells, and commercially, this natural-like RNA duplex species is also known as ‘Larifan’ [69]. Subsequently, the V-dsRNA samples (prepared in 1× PBS) of concentration range 0-2.5 µg mL^-1^ (for B2-ALP, 2.5× dilution at each step) and 0-15 µg mL^-1^ (for B2-NLuc, 3.5× dilution at each step) were added to the plates (100 µL per well). The plates were incubated at 28 °C for 1.5 h for dsRNA binding/capturing to the B2-mScarlet surface. Plates were then washed 3 times by the WB. The next steps for revelation differ depending on the B2-ALP or B2-NLuc scaffold and are discussed in the following paragraphs.

1. Detection by **B2-ALP**: The B2-ALP (0.2 µg mL^-1^) was prepared in conjugate buffer (composition for 1,000 mL, pH 7.4: Tris 2.40 g, NaCl 8 g, PVP K25, M_W_ ∼ 24,000, 20 g, Tween-20 0.5 g, BSA 2 g, MgCl₂·6H₂O 0.2 g, KCl 0.2 g) and added 100 µL per well for revealing the bound dsRNA. The plates were then incubated at 28°C for 30 min and washed with WB for 4 times. Next, the pNPP substrate (1 mg mL^-1^, J&K Scientific GmbH, Cat. 254303) was prepared in substrate buffer (composition for 1,000 mL, pH 9.8: Diethanolamine 105.7 g, MgCl₂·6H₂O 0.02 g) and added 100 µL per well. The plates were incubated in the dark for ∼50-60 min for the color development, and the signal was then measured by the Thermo Fisher Scientific Varioskan microplate reader. The absorbance was recorded at 405 and 492 nm. For the analysis of the data, the 492 nm signal (which refers to the non-specific absorption) and the signal of only the substrate were subtracted from the 405 nm signal, and this was referred to as = Corrected Absorbance.
2. Detection by **B2-NLuc**: B2-NLuc (0.03 µg mL^-1^) was prepared in the buffer: 1× PBS, 0.1% Tween-20, 0.1% BSA, pH 7.4, and 100 µL/well was added for the revelation. The plates were then incubated at 28°C for 30 min and afterward washed with WB for 4 times. The furimazine substrate was dissolved in DMSO and stored as a 2 mg mL^-1^ stock for use. For the assay, the furamizine (4 µg mL^-1^) was prepared in furimazine substrate buffer (composition for 500 mL, pH 6: MES buffer 9.76 g, EDTA 0.2 g, KCl 5.59 g, Thiourea 1.33 g) and added to the plate 100 µL well^-1^. After the addition of the substrate, luminescence was measured (after ∼5 mins) by the Thermo Fisher Scientific Varioskan microplate reader. Both black (Thermo Fisher Cat. 437111) or white (Thermo Fisher Cat. 436110) plates can be used for the B2-NLuc assay; the results reported in this work are obtained by using the white MaxiSorp plate.

Troubleshooting and Quality Control Parameters: **(i)** The washing of the plates in each step should be done carefully. In our assays, we have performed manual hand-held washing using the Thermo Scientific Nunc Immuno Washer (12-channel). Alternatively, an automated plate washer can also be used. **(ii)** NaN_3_ (0.1-0.2 g for 1,000 mL) can be added to the Sand-BIRD assay buffers for longer storage at RT. **(iii)** It is important to note that both the black and white 96-well plates perform near identically for the B2-NLuc assay when normalized signal intensities are compared. However, white plates produce much higher RLU in terms of absolute count and are preferred in this work. **(iv)** Make sure the conjugate and the substate buffer for B2-ALP have MgCl_2_ added in; without this, the ALP enzyme will function sub-optimally since the bacterial ALP requires Mg^2+^ as a cofactor for its activity. **(v)** Using an automated multichannel pipette results in much better reproducibility and consistency in the assay results. We have used the INTEGRA Viaflo Electronic Pipettes for all our experiments. **(vi)** For the B2-NLuc assay, SuperBlock blocking buffer from Thermo Scientific (Cat. 37580) can also be used for the blocking step; this can reduce the blocking step to 15 mins.

### Virus-Infected Plant Samples Preparation

Grapevine fanleaf virus (GFLV isolate GHu) isolate used in this work is described in Vigne et al. (2013) [93]. The Tomato bushy stunt virus (TBSV-BS3Ng) strain used in this work is described in Incarbone et al. (2020) [72] and was obtained from Bioreba AG (Switzerland). Mechanical inoculation of *N. benthamiana* was carried out for the virus infection as per the previously described method by Hull (2009) [94]. The Leaf tissues were processed using the Bioreba-recommended homogenization apparatus. Briefly, 1 g of fresh *Nicotiana Benthamiana* leaves was homogenized in 10 mL (1:10 w/v) of Bioreba Extraction buffer “General” (Composition for 1,000 mL, pH 7.4; 2.4 g Tris, 8 g NaCl, 20 g PVP K25, 0.5 g Tween-20, 0.2 g KCl, 0.2 g NaN₃ per liter, pH adjusted with HCl) using Universal extraction bags (12 × 15 cm; Bioreba, Art. No. 430100) and a Bioreba hand homogenizer with ceramic balls (Art. No. 400011). The crude homogenate was clarified by centrifugation at 14,000 × g for 5 min (at 4 °C), and the supernatant was collected. The clarified supernatant was directly applied to ELISA plates for dsRNA detection by the Sand-BIRD assay, and the response of infected and non-infected samples can be compared. For the extraction of the grapevine samples, the Bioreba Extraction buffer “Grapevine” should be used (Composition for 1,000 mL, pH 8.2; 24 g Tris, 8 g NaCl, 20 g PVP K25,10 g PEG 6000, 0.5 g Tween-20, 0.2 g NaN₃, pH adjusted with HCl). For preparation of virus-infected time course samples, 6-week-old *N. Benthamiana* plants were infected with TBSV (TBSV BS3Ng strain) and GFLV (GFLV GHu strain, which produces visible symptoms in *N. Benthamiana*), and both inoculated and systemic leaves were collected across a span of 20 dpi. The collected leaves were then extracted in Bioreba Extraction buffer “General” (1:10 (w/v) tissue dilution) and analyzed by the Sand-BIRD method. In parallel, identical samples were tested by the Bioreba virus ELISA protocol for TBSV (Art No. 161815 and 161825) and GFLV (Art No. 120412 and 120422) as per the manufacturer’s recommended protocol. The GFLV F13 strain-infected plants were provided by Dr. Olivier Lemaire from INRAE, Colmar, France.

### Mosquito Samples Preparation

*Aedes aegypti* mosquitoes from the BORA colony, originally collected on Bora-Bora Island in 1980 [95], were reared in the VectoPole insectary at MIVEGEC. Eggs were hatched in deionized water, and larvae were fed shrimp pellets (Novo Prawn, JBL) at 26°C under a 12 h:12 h light-dark cycle until pupation. Adults were maintained in BugDorm 4M2222 cages at 28°C and 70% relative humidity under a 14 h:10 h light-dark cycle, with *ad libitum* access to 10% sugar solution. West Nile virus (WNV) strain IS98-ST1, originally isolated from a stork in Israel in 1998 [96], was obtained from the Centre de Ressources Biologiques, Institut Pasteur, Paris. The Dengue virus strain was: DENV D2Y98P strain, 1998 DEN2 Singapore human isolate [97]. The virus was propagated in C6/36 mosquito cells supplemented with 2% FBS, titrated by plaque assay in Vero cells as previously described [98], and stored at −70°C.

Cold-anesthetized female Ae. aegypti mosquitoes (3–5 days old) were intrathoracically injected with 69 nL of RPMI containing 0.5 plaque-forming unit (PFU) of WNV, or 1 or 10 PFU of DENV using a Nanoject III (Drummond Scientific Company) fitted with glass capillary needles (44.45 mm length, 1.14 mm outer diameter; Drummond). Control mosquitoes received the same volume of RPMI alone. Mosquitoes were then maintained under standard rearing conditions for 10 days before being freeze-killed and stored at −70°C. In a separate oral infection assay, 4-day-old female *Ae. aegypti* were starved for 12 h and then offered an infectious blood meal using a Hemotek feeding system (Discovery Workshops) covered with pig intestine. The 2.5 mL artificial blood meal contained one part WNV diluted in RPMI to 2 × 10^7^ PFU mL^-1^, two parts washed rabbit erythrocytes (from animals housed in the BSL2 VectoPole animal facility; authorization no. H3417221), and 25 mM ATP (Thermo Fisher Scientific). Engorged females were selected and maintained under standard rearing conditions for 14 days before analysis.

The successful viral infection in mosquitoes was validated by RT-qPCR. Total RNA was extracted from mosquitoes using the EZNA RNA Extraction Kit I (OMEGA). Positive-strand genomic RNA was absolutely quantified by one-step RT-qPCR in 10 µL reactions containing 5 µL iTaq Universal SYBR Green One-Step Kit (Bio-Rad), 300 nM each forward and reverse primer, as described in Medkour et al., 2025 [99], and 2 µL RNA extract. Amplification was performed on an AriaMx Real-Time PCR System (Agilent) under the following conditions: 50°C for 10 min, 95°C for 1 min, and 40 cycles of 95°C for 10 s and 60°C for 25 s, followed by melt curve analysis. Absolute standard curves for DENV and WNV genomic RNA were generated by amplifying the qPCR target with their respective primers, as previously described [99–101]. The 95% limit of detection (LoD) was determined from six replicate serial dilutions of gRNA templates by calculating the fraction of positive detections [102].

For the analysis of the virus-infected mosquito samples in the Sand-BIRD assay, samples were homogenised in a 1:10 (w/v) tissue-to-buffer ratio (5 mosquitoes in 500 µL of buffer; considering 1 mosquito having an average weight of 10 mg) in ice-cold Bioreba “General” extraction buffer first. Next, the samples were clarified by centrifugation at 14,000 × g for 5 min (at 4°C), the supernatant was collected, and directly subjected to the sandwich assay for dsRNA detection by the B2-ALP sensor scaffold.

### dsRNA Elution for Nanopore Sequencing

For the virus diagnostics using the dsRNA-centred approach, the samples eluted directly from the 96-well microplate were subjected to a downstream Oxford Nanopore sequencing pipeline. The samples were eluted from the 96-well plate after the “sample binding/incubation step” in the following way. After the sample binding (on the 5% NFDM blocked B2-mScarlet surface), the plate was washed with 1× PBS-T (3 times), 100 µL of 2.5 M lithium chloride (LiCl) was added to each well containing positive samples, and shaking of the microplate was carried out for 30 mins at 400 RPM (in RT). Next, a total of 6 wells were pulled in a 1.5 mL microcentrifuge tube, and 30 µg of GlycoBlue (Thermo Fisher, Cat. AM9515) was added; the tube was kept on ice for 2-3 min, and an equal volume of 600 µL cold isopropanol (100%) was added. The tube was then inverted 4-5 times, and kept in -80°C for 15 mins. A centrifugation was carried out next (18,000 ×*g*, 5 mins, at 4°C) and everything except the pellet was removed by pipetting slowly. The pellet was then washed with 70% ethanol (500 µL) and centrifuged again (18,000 ×*g*, 5 mins, at 4°C). The entire ethanol solution was discarded slowly, the tube was kept at 37°C for 5 min for drying, and finally, the pellet was dissolved in 8 µL of DNase/RNase-free water (or autoclaved MilliQ water). This sample is essentially the nucleic acid species captured by the B2 surface, which can be subjected to nanopore library preparation either immediately or can be stored at -20°C.

### Nanopore Direct dsRNA Sequencing Pipeline (dsNanoSeq)

As a first step of the nanopore sequencing library preparation, the dsRNA samples eluted from the plate were polyadenylated. For the polyadenylation reaction, following components were added (in a 200 µL tube) to a total volume of 10 µL: 6 µL of dsRNA sample, 1.25 µL of DNase/RNase free water, 0.25 µL of RNase inhibitor (Promega, Cat. N2611), ATP 1 µL (from 10 mM stock, NEB, Cat. P0756S), 1 µL of Poly(A) Polymerase Reaction Buffer (NEB, Cat. M0276S), 0.5 µL of *E. coli* Poly(A) Polymerase (NEB, Cat. M0276S). The tube was then immediately placed in a thermocycler at 37°C for 5 mins, heated to 70°C (5 mins), and placed on ice.

The nanopore sequencing was carried out by the direct RNA sequencing (SQK-RNA004) kit as per the recommended protocol by Oxford Nanopore. In our experiment, we ligated a custom RT adaptor containing a barcode to the polyadenylated dsRNA species for demultiplexing of multiple samples. However, the RT step was skipped owing to the fact that the dsRNA is essentially a thermodynamically stable duplex structure. For the details of this custom RT adaptor (WDX_RTA), refer to the original WarpDemuX article [70] and GitHub repository; https://github.com/KleistLab/WarpDemuX. These WDX_RTA adaptors were kindly provided by the Redmond Smyth lab from Institut de biologie moléculaire et cellulaire (IBMC). For the ligation reaction, a total of 15 µL reaction mixture is prepared with the following components: 8.5 µL of polyadenylated dsRNA from the previous step, 3 µL NEBNext Quick Ligation Reaction Buffer (NEB, Cat. B6058), 1 µL of RNase inhibitor (Promega, Cat. N2611), 1 µL of custom WDX RT adaptor (WDX_RTA), 1.5 µL T4 DNA Ligase 2M U mL^-1^ (NEB, Cat. M0202M). For each sample, one specific barcode was used; see the following section for barcode details. Next, mix all the components by pipetting, quick spin down (on a tabletop centrifuge), and incubate at room temperature for 10 min (first ligation reaction step). The stock of the Agencourt RNAClean XP (Beckman Coulter) beads was resuspended by vortexing, and 15 µL of beads was added to a 1.5 mL LoBind tube, and the ligation reaction mixture was added to this. After mixing by pipetting, the tubes are kept on a Hula rotator mixer (or on a standing rack with occasional tapping on the tube) for 5 min at room temperature. The samples were cleaned subsequently as per the ONT DRS_9195_v4_revF_11Dec2024 protocol (https://nanoporetech.com/document/direct-rna-sequencing-sqk-rna004). Briefly, the adaptor ligated dsRNA samples are captured by Agencourt RNAClean XP beads (Beckman Coulter, A63987); next, the beads are pelleted down on a magnetic rack and washed by 70% freshly prepared ethanol, and finally, the dsRNA bound to the beads was eluted with 23 µL nuclease-free water. Next, all the individual samples for each barcode were pooled together, and ligation of the RNA Ligation Adapter (RLA) was performed as per the ONT protocol. Cleaning of the dsRNA samples was done next to remove the unbound RLA (Agencourt RNAClean XP beads approach as mentioned earlier), the final library was eluted in 13 µL RNA Elution Buffer (REB), and the sample (1 µL) was quantified using the Qubit fluorometer using the Qubit dsDNA HS assay (Invitrogen, Cat. Q32851).

For the control, Larifan (*bona fide* dsRNA) and 142 bp dsRNA, we used MinION flow cell. And for the TBSV, GFLV, and WNV samples analysis, we employed PromethION cells from Oxford nanopore and performed a run of approximately 24 hrs for each case. The library prepared in the previous step was loaded in the flow cell by strictly following the ONT protocol guidelines for MinION and PromethION flow cells. We ran a total of 3 independent sequencing runs. Run1 (in MinION flow cell): Control dsRNA samples – WDX_RTA_BC5: 142 bp dsRNA, WDX_RTA_BC7: Larifan, *bona fide* dsRNA. Run2 (in PromethION flow cell): virus-infected plant samples eluted from 96-well plate - WDX_RTA_BC3 and 4: TBSV, WDX_RTA_BC5 and 6: GFLV. Run3 (in PromethION flow cell): virus-infected mosquito samples eluted from 96-well plate - WDX_RTA_BC3 and 4: WNV. Larifan and the 142 bp dsRNA control samples were directly subjected to poly(A) tailing and nanopore library preparation as obtained (without any elution from the microplate step). This run (Run 1) was performed first to validate the direct nanopore dsRNA sequencing workflow (dsNanoSeq). Afterwards, in Runs 2 and 3, samples were first eluted from the plate and then sequenced to validate the integrated sensing-to-sequencing BIRD-Seq workflow for virus identification.

### dsNanoSeq Data Analysis

Raw nanopore data from the experiment stored in POD5 format were basecalled using Dorado v1.0.2 (Oxford Nanopore Technologies) with a high accuracy (HAC) model, employing GPU acceleration (NVIDIA CUDA) on the France IFB Core Cluster. The resulting output was a basecalled file in BAM format, which was used for subsequent alignment and mapping steps using a custom-written modular bash pipeline. Quality filtering was conducted on the basecalled BAM file with Chopper (v0.9.0) [103], using a Phred quality score (Q) cutoff of 10. Barcode demultiplexing was achieved using the WarpDemuX algorithm [70], retaining only reads with a ≥0.98 prediction confidence scores to minimise misassignment and mapping to high-quality Q10 reads. Taxonomic classification of the filtered reads was done using Kraken2 (v2.1.3) [104] against a locally indexed viral RefSeq nucleotide database. To minimise the false-positive assignments, all the Kraken2 candidate species were subjected to a genome coverage validation step: reads were mapped to the corresponding reference sequence using minimap2 (map-ont preset; k-10, v2.28), and only the species achieving >20% genome breadth coverage was considered as true confirmed hits. The reads are mapped to the reference genome of the specific virus species (TBSV – Tomato bushy stunt virus *Arabidopsis* strain BS3Ng; GFLV - Grapevine fanleaf virus isolate GHu, RNA1 - JN391442.1, RNA2 - EF426852.1; WNV - West Nile virus lineage 1, NC_009942.1) to decipher the mapping statistics including forward/reverse strand coverage, mean depth, and genome coverage (calculated by SAMtools, v1.21). The host reads for Larifan were found by mapping the reads to the *Escherichia coli* K-12 strain genome (Accession: NC_000913.3). For mapping of the plant host *Nicotiana benthamiana*, the NbKLAB assembly was used (Accession: JAXGFW00000000041) [105]. The host reads for WNV were mapped to the *Aedes aegypti* whole genome (Accession: NC_035107.1). In order to recover the maximum number of reads that map to the target virus reference, a series of steps was done sequentially: minimap2 (k=10 criteria), a BLASTn (e-value ≤ 1×10^-5^), minimap2 (k=7 criteria), and a final BLASTn (e-value ≤ 1×10^-3^). The reads distribution was visualized by NanoPlot (v1.46.2) [103]. All the plots were generated either by custom-written R scripts or by OriginLab Origin 2026.

## Supporting information

The supplemental information contains (1) AlphaFold modelling of protein constructs, (2) Binding kinetics from SPR, (3) Illustration of dot blot.

## Resource Availability

### Lead contact

Requests for further information and resources should be directed to and will be fulfilled by the lead contact, Christophe Ritzenthaler.

### Data and code availability

All data reported in this paper will be shared by the lead contact upon request. Any additional information required to reanalyse the data reported in this paper is available from the lead contact upon reasonable request.

### Author contributions

SSzunerits, TB, CR: Conceptualization

ANX, SS, HS, EFM, AA, DC, CR: Methodology, Formal analysis

ANX, SS: Investigation, Validation, Data Curation

ANX, SS, HS, TB, CR: Writing - Original Draft

EFM, DC, BM, JP, SSzunerits, TB, CR: Writing - Review & Editing

ANX, SS, TB: Visualization, Software

SS, CR: Supervision, Project administration

JP, SSzunerits, TB, CR: Funding acquisition

## Acknowledgements

We thank Dr. Dace Pjanova (Riga Stradins University, Latvia) for kindly providing the viral dsRNA (Larifan). We are grateful to Bruno Müller for the kind gift of the pLip6v plasmid. We also acknowledge Dr. Olivier Lemaire (INRAE, Colmar, France) for providing the GFLV-F13 strain-infected plants, and we thank the Redmond Smyth lab at the Institut de biologie moléculaire et cellulaire (IBMC) for providing the custom WDX_RTA adaptors used for the nanopore direct dsRNA sequencing pipeline.

## Supplemental information

Supplemental information related to this article can be found online.

The supplemental information contains (1) AlphaFold-3 modelling of protein constructs, (2) Binding kinetics from SPR experiment, (3) Schematic illustration of dot blot experiment, (4) data related to nanopore sequencing.

## Assay and Pipeline Nomenclature

- **Res-BIRD**: Surface plasmon resonance (SPR) platform for characterizing protein-dsRNA binding kinetics.
- **Dot-BIRD**: B2-Integrated dsRNA Detection dot blot assay.
- **Sand-BIRD**: B2-Integrated dsRNA Detection sandwich assay.
- **dsNanoSeq**: Nanopore direct dsRNA sequencing methodology for the identification of B2-captured dsRNA.
- **BIRD-Seq**: B2-Integrated dsRNA Detection & Sequencing pipeline. The whole end-to-end agnostic pipeline of dsRNA detection through Sand-BIRD and the subsequent nanopore sequencing pipeline (dsNanoSeq) is referred to as BIRD-Seq.

## Glossary and Abbreviations

Sand-BIRD: B2-Integrated dsRNA detection sandwich assay. This refers to the B2-based assay for the dsRNA monitoring, wherein B2-mScarlet (B2 fused to mScarlet) is used as a dsRNA capturing interface and either B2-ALP (B2 fused to alkaline phosphatases) or B2-NLuc (B2 fused to NanoLuc luciferase) for detecting the bound dsRNA.
BIRD-Seq: B2-Integrated dsRNA detection & sequencing pipeline. The whole end-to-end agnostic pipeline of RNA duplex detection through Sand-BIRD and the subsequent nanopore direct dsRNA sequencing pipeline (dsNanoSeq) is referred to as BIRD-Seq.

