## Supplementary material for "BIRD-Seq: B2 Protein Integrated End-to-End Pipeline for dsRNA Detection and Nanopore Sequencing for Virus Monitoring": The supplemental information contains (1) AlphaFold modelling of protein constructs, (2) Binding kinetics from SPR, (3) Illustration of dot blot.

### Supporting Figure/Table

Table S1. Sequence information of all the protein constructs used in this work.

| |
|---|
| B2-SpyTag003 (B2-SpT), 11.8 kDa<br>MPSKLALIQELPDRIQTAVEAAMGMSYQDAPNNVRRDLDNLHACLNKAKLTVSRMVTSLLEKPSVVAYL<br>EGKASGGGSGGGSGGGRGVPHIVMVDAYKRYKSGGGHHHHHH |
| Cys-SpyCatcher003 (C-SpC), 13.6 kDa<br>MHHHHHHCGGGSGGGVTTLSGLSGEQGPSGDMTTEEDSATHIKFSKRDEDGRELAGATMELRDSSGKTISTWI<br>SDGHVKDFYLYPGKYTFVETAAPDGYEVATPIEFTVNEDGQVTVDGEATEGDAHT |
| B2-mScarlet, 35.8 kDa<br>MAGMPSKLALIQELPDRIQTAVEAAMGMSYQDAPNNVRRDLDNLHACLNKAKLTVSRMVTSLLEKPSVV<br>AYLEGKAYSGGGGVSKGEAVIKEFMRFKVHMEGSMNGHEFEIEGEGEGRPYEGTQTAKLKVTKGGPLPF<br>SWDILSPQFMYGSRAFIKHPADIPDYYKQSFPEGFKWERVMNFEDGGAVTVTQDTSLEDGTLIYKVKLR<br>GTNFPPDGPVMQKKTMGWEASTERLYPEDGVLKGDIKMALRLKDGGRYLADFKTTYKAKKPVQMPGAYN<br>VDRKLDITSHNEDYTVVEQYERSEGRHSTGGMDELYKHHHHHH |
| B2-ALP, 56.5 kDa<br>MPSKLALIQELPDRIQTAVEAAMGMSYQDAPNNVRRDLDNLHACLNKAKLTVSRMVTSLLEKPSVVAYL<br>EGKAGGGSRTPEMPVLENRAAQGDITAPGGARRLTGDQTAALRDSLSDKPAKNIILLIGDGMGDSEITA<br>ARNYAEGAGGFFKGIDALPLTGQYTHYALNKKTGKPDYVTDSAASATAWSTGVKTYNGALGVDIHEKDH<br>PTILEMAKAAGLATGNVSTAELQGATPAALVAHVTSRKCYGPSATSEKCPGNALEKGGKGSITEQLLNA<br>RADVTLGGGAKTFAETATAGEWQGKTLREQAQARGYQLVSDAASLNSVTEANQQKPLLGLFADGNMPVR<br>WLGPKATYHGNIDKPAVTCTPNPQRNDSVPTLAQMTDKAIELLSKNEKGFFLQVEGASIDKQNHAANPC<br>GQIGETVDLDEAVQRALEFAKKEGNTLVIVTADHAHASQIVAPDTKAPGLTQALNTKDGAVMVMSYGNS<br>EEDSQEHTGSQLRIAAYGPHAANVVGLTDQTDLFYTMKAALGLKLEHHHHHH |

B2-NLuc, 28.5 kDa

```
MAGMPSKLALIQELPDRIQTAVEAAMGMSYQDAPNNVRRDLDNLHACLNKAKLTVSRMVTSLLEKPSVV
AYLEGKAYSMVFTLEDFVGDWRQTAGYNLDQVLEQGGVSSLFQNLGVSVTPIQRIVLSGENGLKIDIHV
IIPYEGLSGDQMGQIEKIFKVVYPVDDHHFKVILHYGTLVIDGVTPNMIDYFGRPYEGIAVFDGKKITV
TGTLWNGNKIIDERLINPDGSLLFRVTINGVTGWRLCERILASHHHHHH
```

Table S2. SPR Binding affinities and kinetic parameters of various bioreceptors analysed in this work are provided. The affinity was monitored for dsRNA of natural origin (V-dsRNA), and all the $k_{on}$, $k_{off}$, and $K_D$ values are listed.

| **Protein Construct** | **$K_D$ (M)** | **$k_{on}$ ($M^{-1}$ $s^{-1}$)** | **$k_{off}$ ($s^{-1}$)** |
|---|---|---|---|
| *Kinetic Parameters for bona fide viral dsRNA (V-dsRNA)* | | | |
| **B2-SpT** | $8.6 \pm 2 \times 10^{-9}$ | $8.7 \pm 2 \times 10^{4}$ | $7.2 \pm 0.1 \times 10^{-4}$ |
| **B2-mScarlet** | $28 \pm 2 \times 10^{-9}$ | $9.8 \pm 0.8 \times 10^{4}$ | $2.8 \pm 0.2 \times 10^{-3}$ |
| **B2-NLuc** | $54 \pm 14 \times 10^{-9}$ | $4.6 \pm 1.8 \times 10^{4}$ | $2.4 \pm 0.1 \times 10^{-3}$ |
| **B2-ALP** | $105 \pm 16 \times 10^{-9}$ | $1.5 \pm 0.3 \times 10^{4}$ | $1.5 \pm 0.1 \times 10^{-3}$ |
| **J2 Antibody** | $6.4 \pm 1.3 \times 10^{-9}$ | $3.7 \pm 0.4 \times 10^{4}$ | $2.4 \pm 0.2 \times 10^{-4}$ |

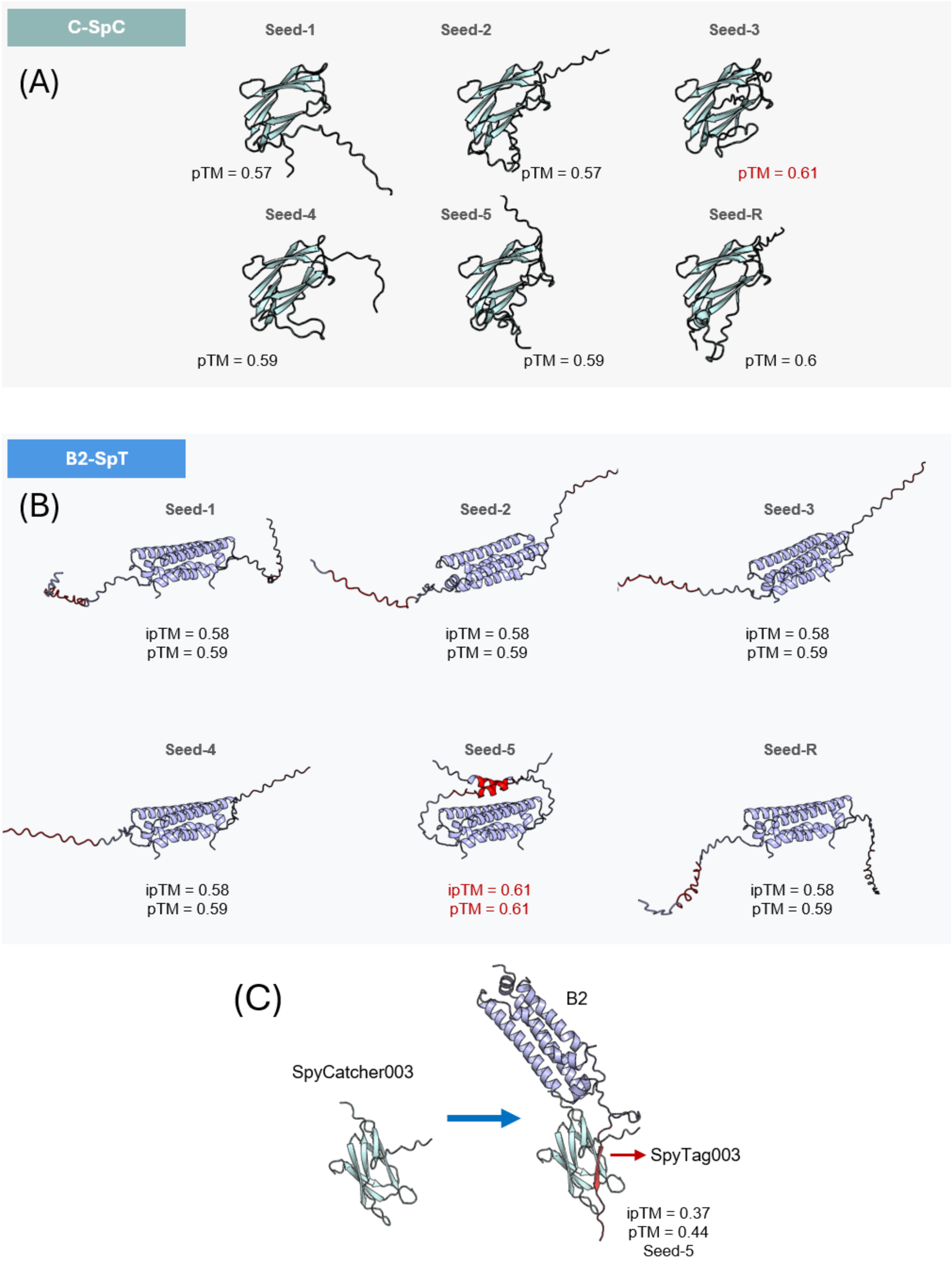


Figure S1. AlphaFold-3 modelling of the (**A**) Cysteine SpyCatcher003 (C-SpC) and (**B**) B2-SpyTag003 (B2-SpT) protein construct. Structures obtained using 5 different seeds (Seed-1 to -5) and a random seed (Seed-R) are shown with respective pTM (predicted Template Modeling score) and ipTM (interface predicted Template Modeling) scores. The score from the best seed is highlighted in red. (**C**) B2-SpT interaction with the C-SpC scaffold as predicted by the AlphaFold-3. The SpyTag003 part of the B2 protein spontaneously interacts with its partner SpyCatcher003 scaffold *in vitro* and corroborates the established SpyTag003/ SpyCatcher003 interaction chemistry. This immobilization process here was carried out over a gold prism, within the SPR instrument (follow Figure 2A).

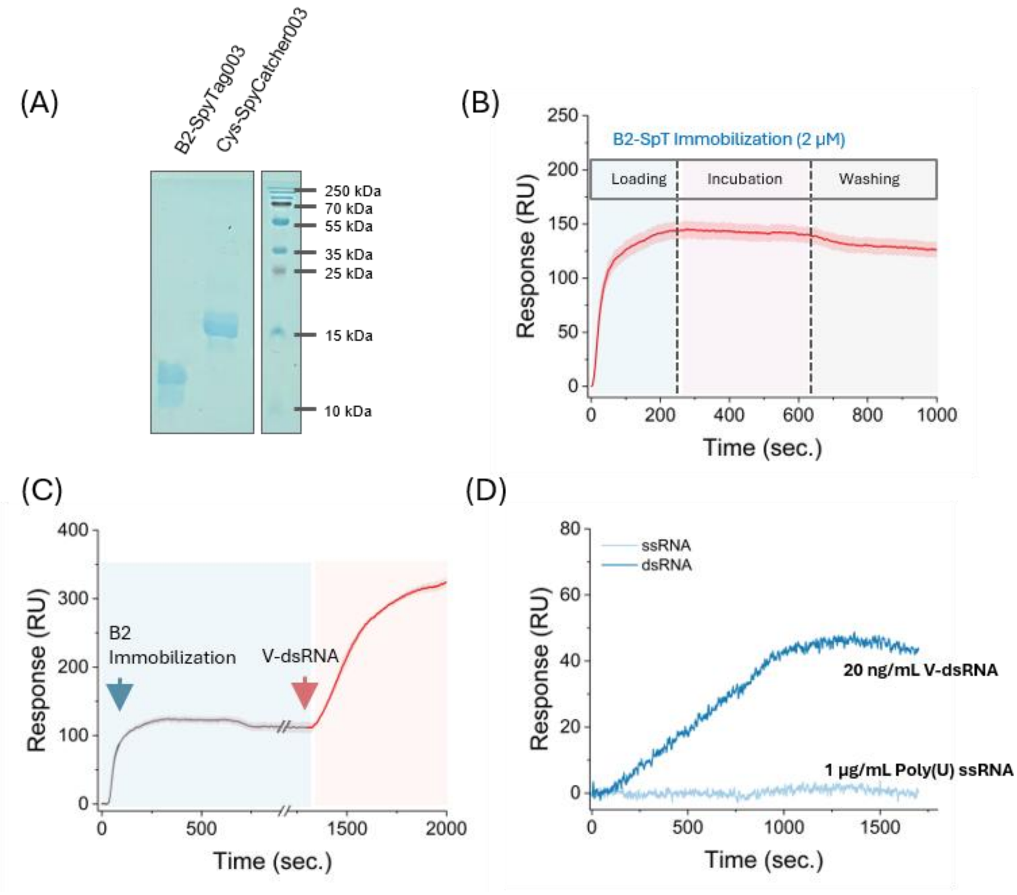


Figure S2. (**A**) SDS-PAGE gel of the protein construct B2-SpyTag003 and Cys-SpyCatcher003 stained by Coomassie blue. (**B**) RU response of the B2-SpyTag003 (B2-SpT) immobilization over the Cys-SpyCatcher003 surface (schematically shown in Figure 2A, step-2). The signal rises upon introduction of the B2-SpT, and stabilizes around 125 RU after washing, suggesting successful capturing of the B2 protein over the surface. (**C**) Viral dsRNA (V-dsRNA) injection over the B2-SpT immobilized surface. The V-dsRNA interacts and binds to the surface captured B2, which in turn further increases the RU signal. (**D**) Selectivity profile of B2 bioreceptor in the SPR setup for dsRNA and Poly(U) ssRNA. B2 does not produce any observable RU signal for ssRNA species, even for a 50x higher concentration.

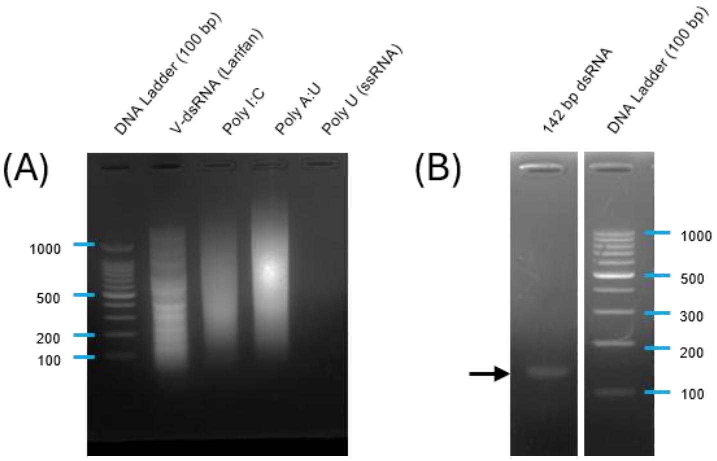


Figure S3. Agarose gel (2%) image of different dsRNA species used in this work. (**A**) V-dsRNA, Poly(I:C), Poly(A:U), and Poly(U) ssRNA and (**B**) 142 bp dsRNA positive control (from Jena Biosciences) are shown. 2 µg of every dsRNA sample is loaded and visualized through ethidium bromide (EtBr) staining. The ssRNA species is not visible in the EtBr-stained gel.

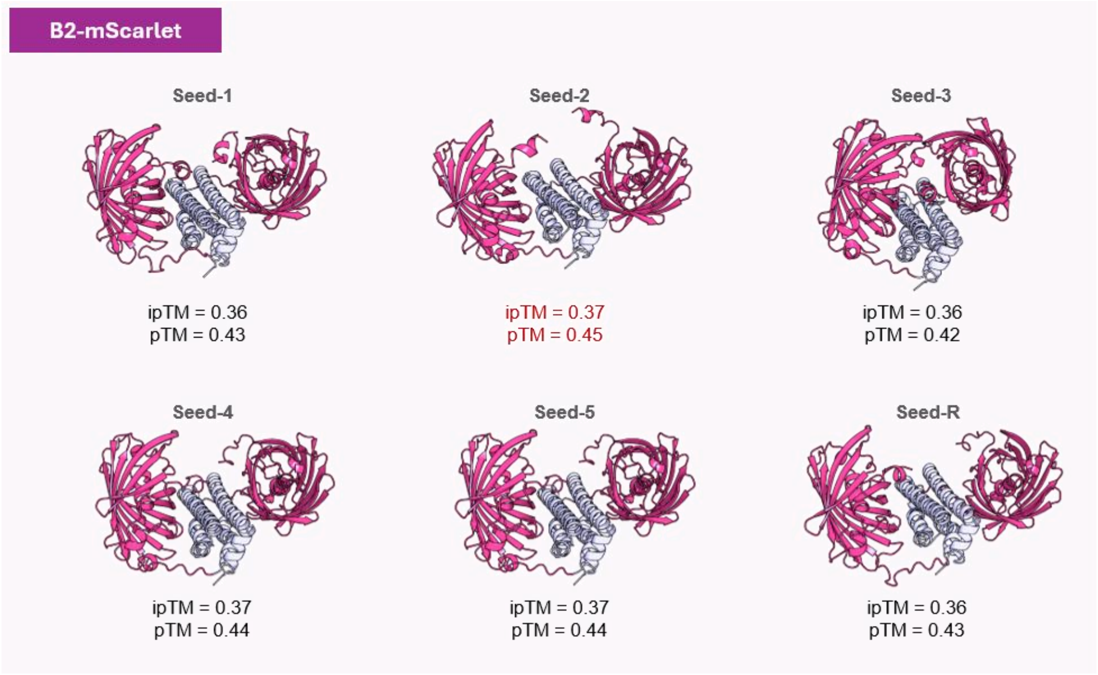


Figure S4. AlphaFold-3 predicted models of B2 fused to mScarlet (B2-mScarlet). The B2 and mScarlet domains are shown in light blue and magenta colors. Structures obtained using 5 different seeds and a random seed are shown with respective pTM (predicted Template Modeling score) and ipTM (interface predicted Template Modeling) scores. The best score from seed-2 is highlighted in red.

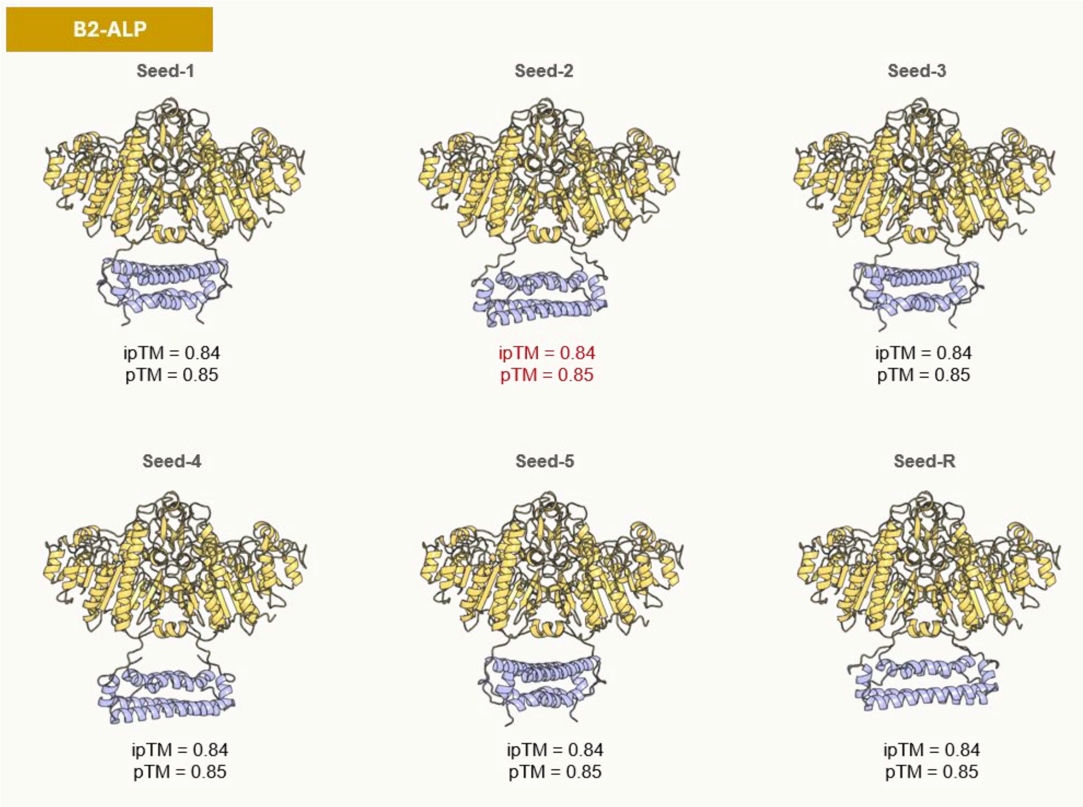


Figure S5. AlphaFold-3 generated models of B2 fused to alkaline phosphatase (B2-ALP). The B2 and ALP domains are shown in light blue and yellow. Structures obtained using 5 different seeds and a random seed are shown with respective pTM (predicted Template Modeling score) and ipTM (interface predicted Template Modeling) scores. In the case of B2-ALP, all the models were predicted to have identical pTM and ipTM values with almost identical spatial orientation of the domains.

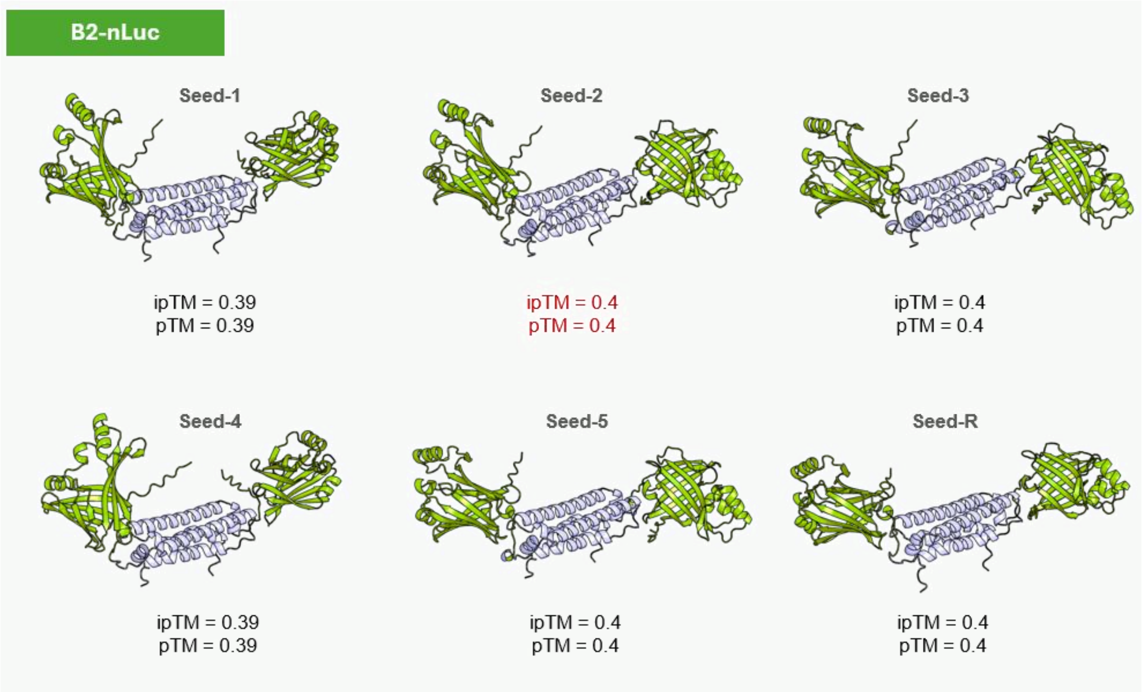


Figure S6. Structure of B2 fused NanoLuc luciferase (B2-NLuc) as predicted by AlphaFold-3. The domains of B2 and NLuc are highlighted by light blue and green colors. Structures obtained using 5 different seeds and a random seed are shown with respective pTM (predicted Template Modeling score) and ipTM (interface predicted Template Modeling) scores. The best score from seed-2 is shown in red.

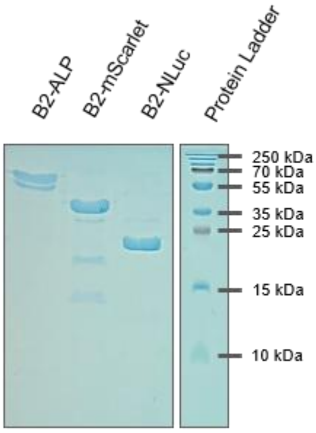


Figure S7. SDS-PAGE electrophoresis gel image of the B2 fusion protein constructs (B2-ALP, B2-mScarlet, B2-NLuc) used for Dot-BIRD and Sand-BIRD dsRNA detection methods. All gel was stained by Coomassie Brilliant Blue-based staining.

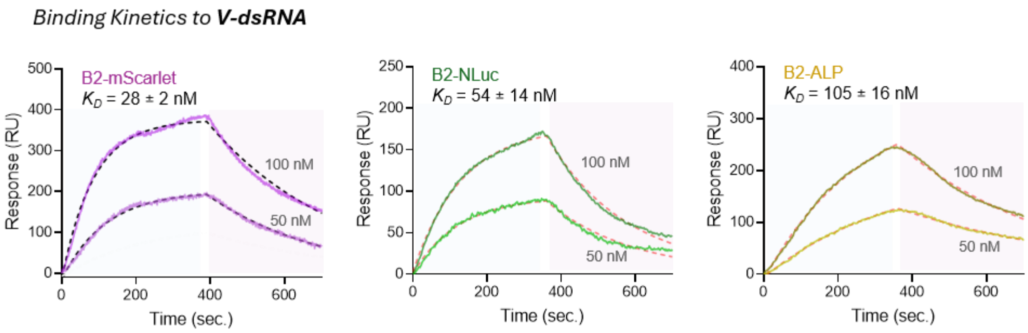


Figure S8. SPR Binding kinetics studies of B2-mScarlet, B2-NLuc, and B2-ALP constructs. All the protein constructs are tested against the *bona fide* viral dsRNA (V-dsRNA), revealing a binding affinity ($K_D$) in the order of nanomolar range.

Dot-BIRD: B2-Integrated Dot Blot for dsRNA Detection

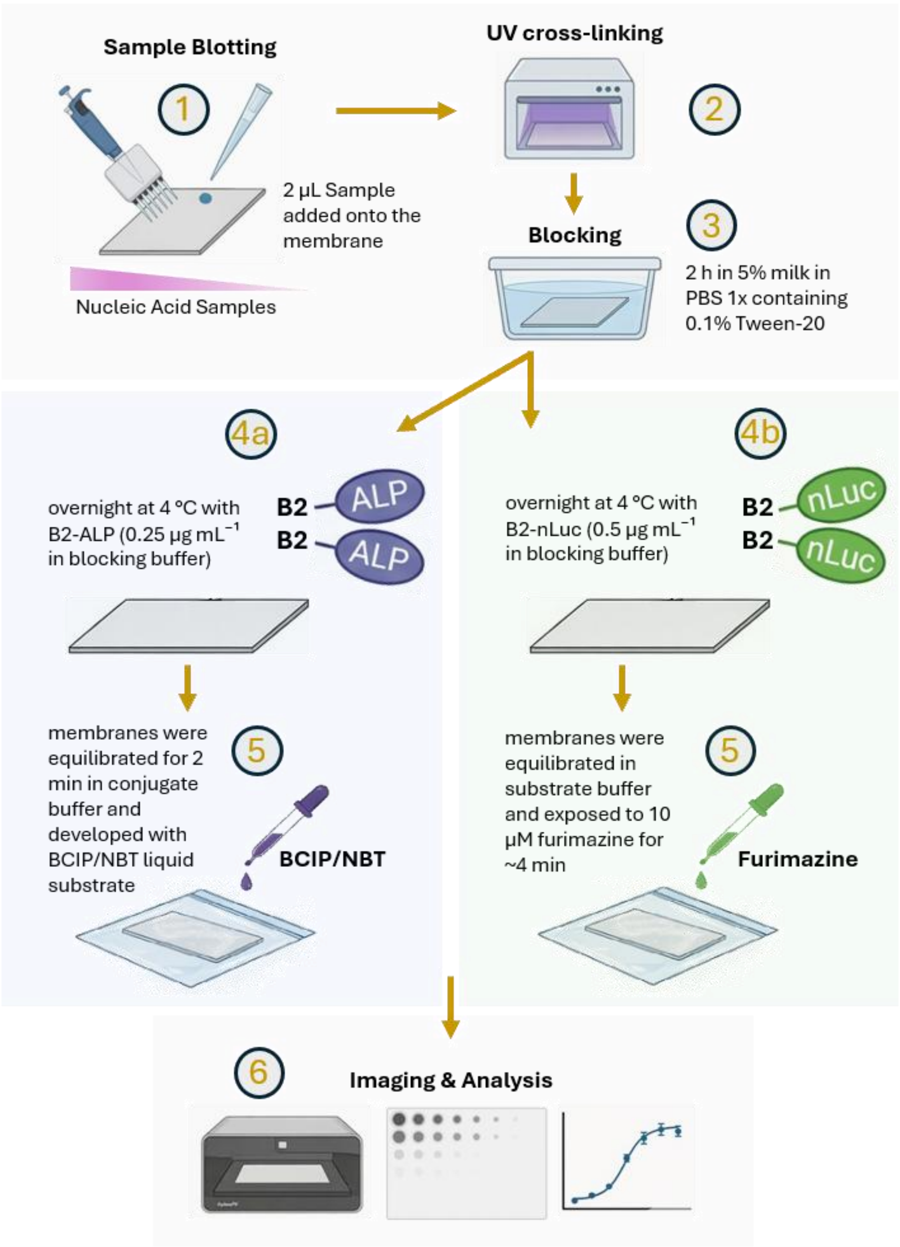


Figure S9. Schematic representation of the B2-Integrated dot blot for dsRNA detection assay (Dot-BIRD). The nucleic acid samples are blotted onto a membrane, UV crosslinked, and blocked first. Then, either B2-ALP or B2-NLuc reporter module is incubated with the samples, which was further revealed by their corresponding substrates (either BCIP/NBT or furimazine, respectively) (see materials and methods for details).

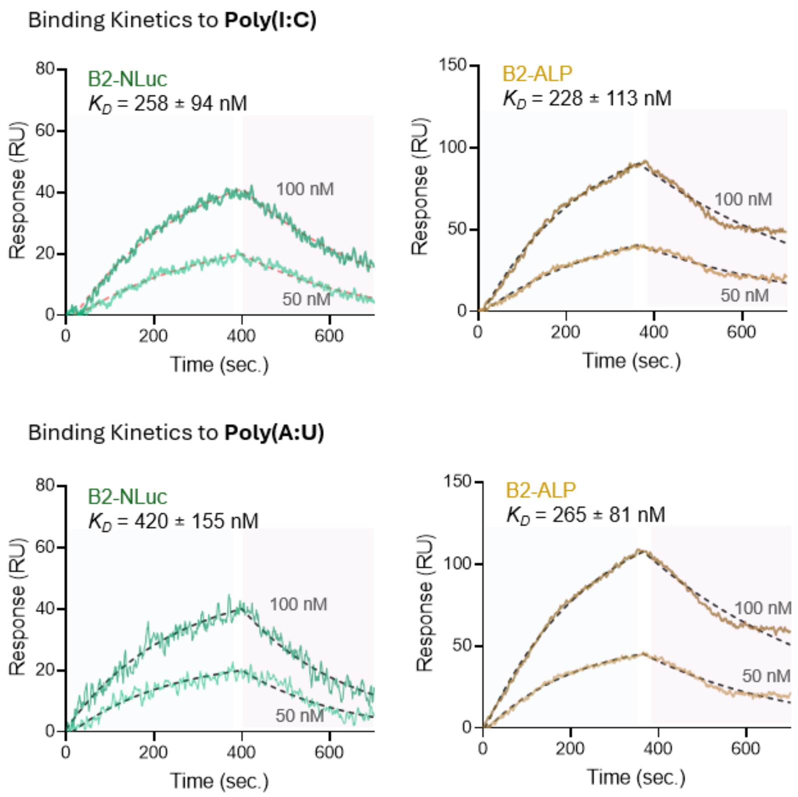


Figure S10. Binding kinetics studies of B2-NLuc and B2-ALP fusion reporter proteins for Poly(I:C) and Poly(A:U) dsRNA species. The B2 bioreceptor revealed an inherent weaker affinity towards synthetic dsRNA species such as Poly(I:C) and Poly(A:U) (as also seen in the Dot blot), and the $K_D$ reduces almost 2-fold in comparison to the natural viral dsRNA (see Figure S8).

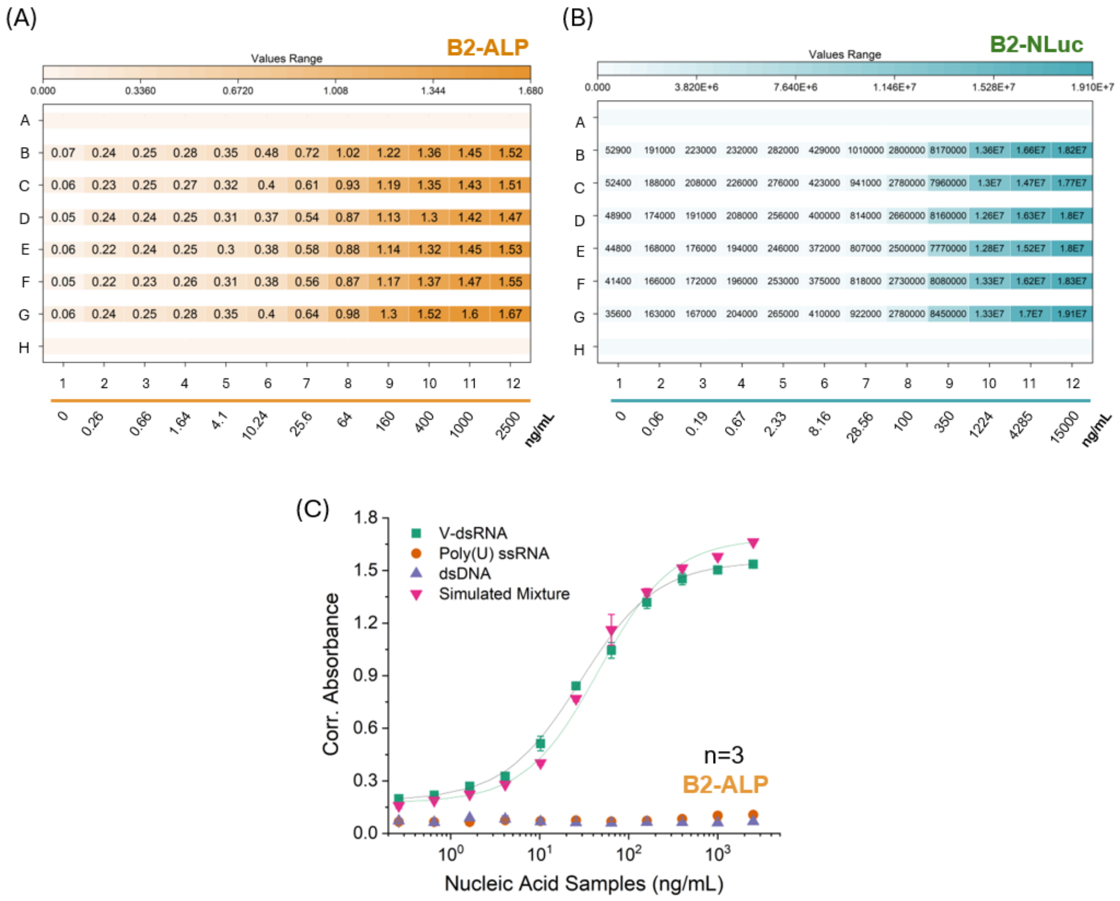


Figure S11. Representative raw values for both the (**A**) B2-ALP and (**B**) B2-NLuc Sand-BIRD assay method for dsRNA detection. For the B2-ALP reporter system, the values are corrected absorbance, and for the B2-NLuc system, RLU values are shown. 6 technical replicates (n=6) are shown for each case when tested against the V-dsRNA. The corresponding nucleic acid concentrations of the standard curve are shown on the x-axis for both the B2 constructs. (**C**) Selectivity profile of the B2-ALP Sand-BIRD assay. It produces no signal for both ssRNA and dsRNA in comparison to the V-dsRNA. The signal response is nearly identical for pure V-dsRNA and a simulated mixture of V-dsRNA, ssRNA, and dsDNA of equal concentration.

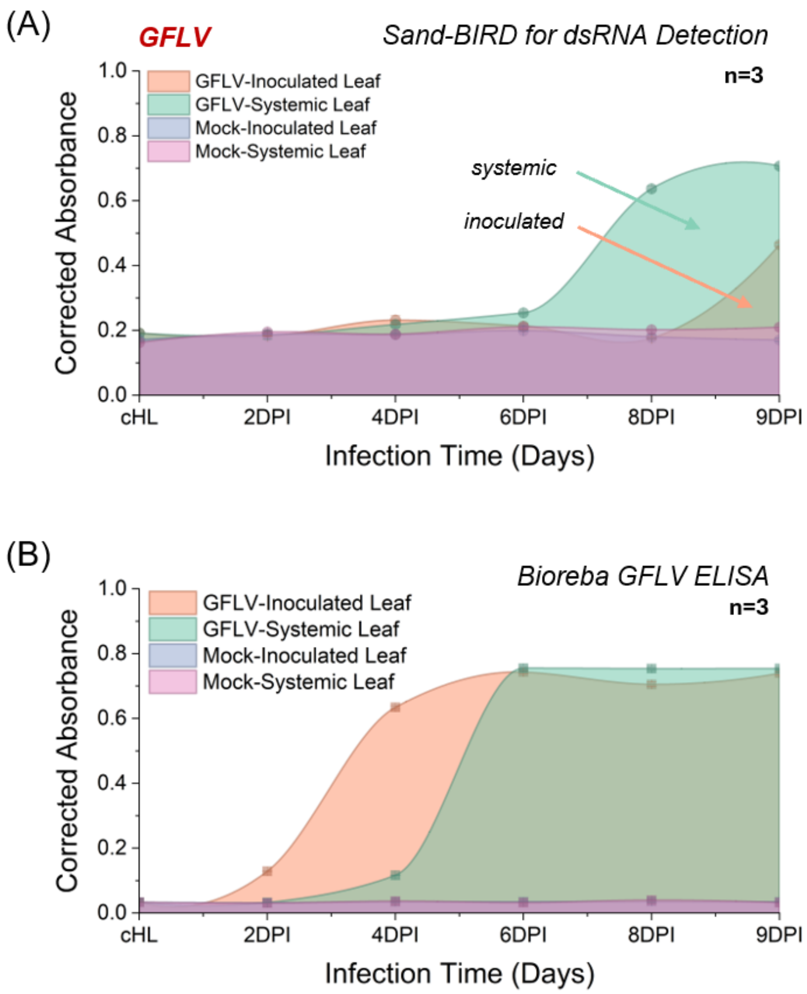


Figure S12. Time course assay for monitoring dsRNA accumulation and GFLV infection in *Nicotiana benthamiana* plants. (**A**) The B2-ALP sandwich assay detects dsRNA in both systemic and inoculated leaves starting from around 7DPI and 9DPI, respectively. The presence of the virus was also confirmed by the (**B**) Bioreba GFLV ELISA in both inoculated and systemic leaves from 4DPI onwards. Response profiles of the mock samples were shown in each case as control samples. The B2 assay monitors actively replicating virus through the presence of dsRNA, whereas the Bioreba virus ELISA detects the virus particle population.

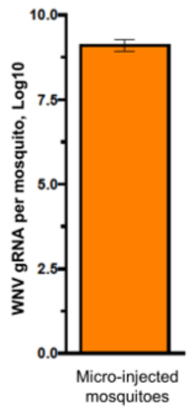


Figure S13. Validation of WNV infection levels in mosquitoes by RT-qPCR. gRNA copies per mosquito that were micro-injected with 0.5 plaque-forming unit (PFU) of WNV. Mosquitoes were collected at 10 days post micro-injection. Bars show geometric mean ± 95% of confidence interval (C.I.); sample size (n) = 30. The healthy control mosquito samples do not produce any signal (data not shown).

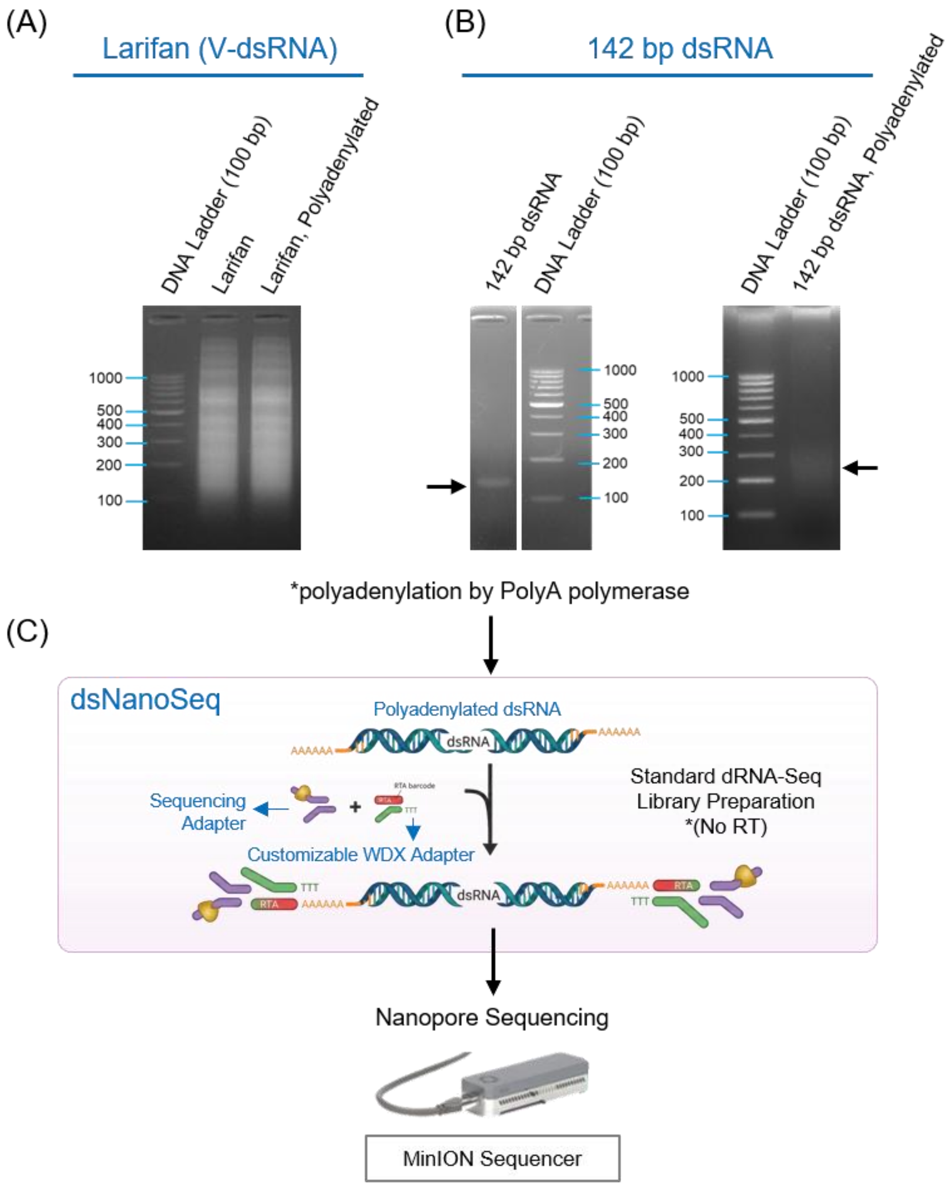


Figure S14. Agarose gel image (2%) showing the polyadenylation of (**A**) Larifan and (**B**) 142 bp dsRNA positive control sample. The gel was stained with Ethidium bromide (EtBr). The polyA tailing was carried out as a first step towards Oxford Nanopore direct RNA sequencing (dRNA-seq) library preparation. (**C**) The next steps of dRNA-seq are shown schematically. A customizable adaptor containing the barcode (from WarpDemuX demultiplexing tool; see methods for adaptor specifications) was ligated to the polyadenylated dsRNA before ligating the sequencing/motor protein adaptor. Both the ligation steps involve cleaning the dsRNA species on magnetic beads to remove excess adaptors. This approach is optimized for dsRNA species and does not involve any RT step (referred to as dsNanoSeq: Nanopore direct dsRNA sequencing). The final library containing both the Larifan and 142 bp dsRNA was sequenced by a MinION sequencer.

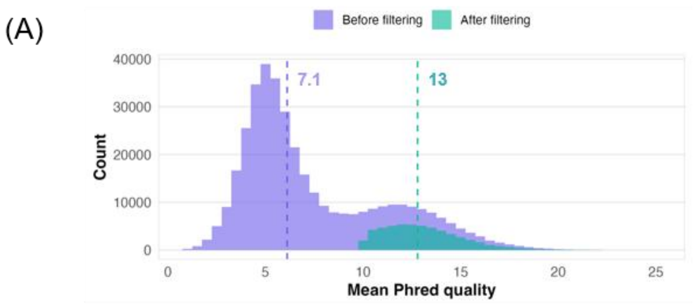


| Stage | | Reads | % |
|---|---|---|---|
| Original extracted reads | : | 380989 | 100% |
| After Chopper QC | : | 54135 | 14.2% |

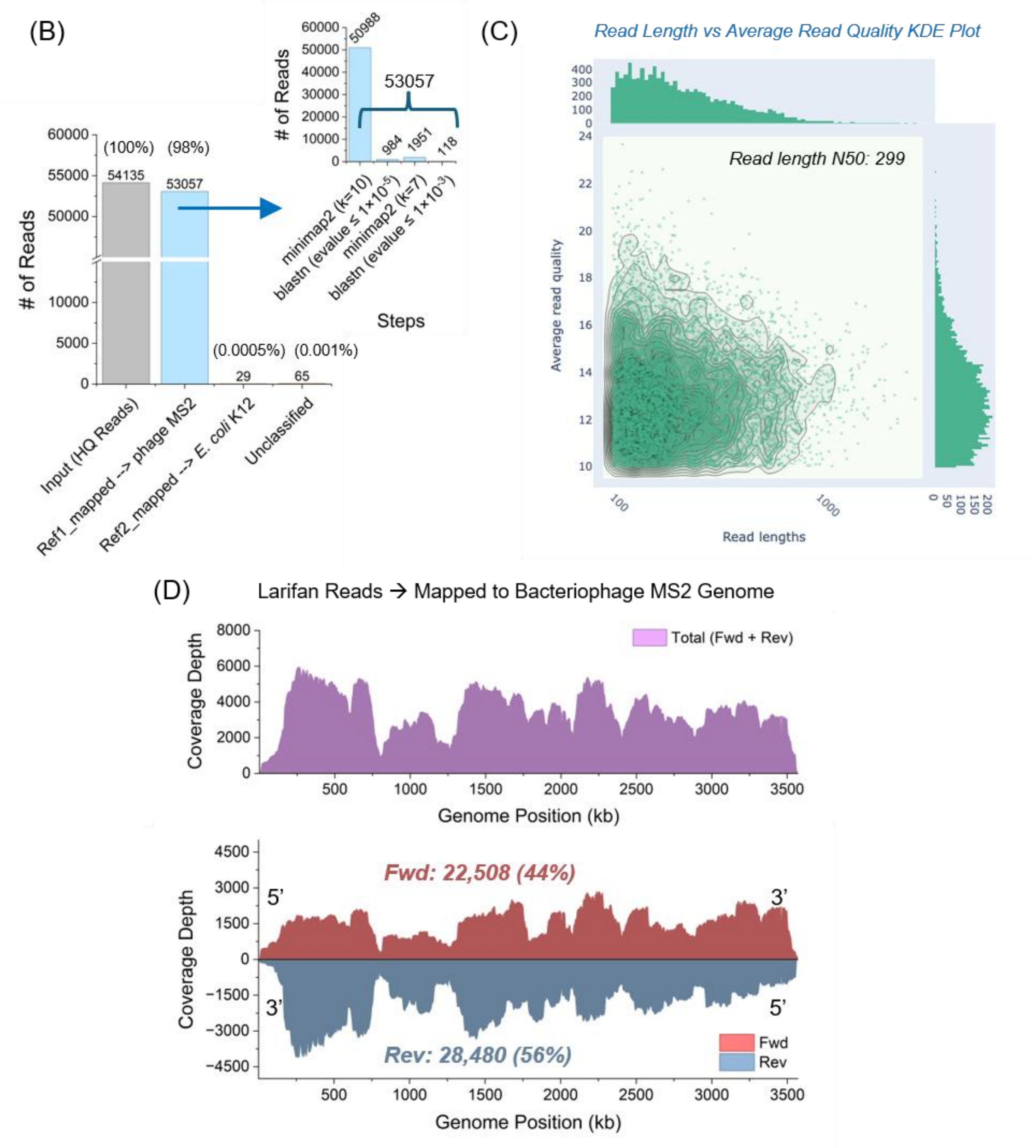


Figure S15. Nanopore sequencing of *bona fide* viral dsRNA species (V-dsRNA, also referred to as Larifan). (**A**) Distribution of the reads obtained from the 'Larifan' sample barcode (WDX_RTA_BC7) in terms of Phred quality. The distribution before and after Chopper quality control (QC) filtering (Phred score (*Q*) >10) is shown with the mean Q score (Q=7.1, before filtering; Q=13, after filtering). Around 14% of high-quality (HQ) reads from the entire barcode population (54,135 reads) were used for further analysis. (**B**) Statistics of the HQ reads mapping to the virus reference genome (bacteriophage MS2 in this case) and host (*E. coli* in this case). Around 98% of all reads map to the reference genome of the virus. In order to recover the maximum number of reads that map to the reference, a series of steps was done sequentially: minimap2 (k=10 criteria), a BLASTn (e-value $\leq 1\times10^{-5}$), minimap2 (k=7 criteria), and a final BLASTn (e-value $\leq 1\times10^{-3}$). Each step recovers a certain number of reads totalling 53,057 reads. (**C**) Distribution of read length vs. average read quality obtained from the NanoPlot package. The Larifan reads population is biased towards shorter reads with a N50 of 299 nt. (**D**) The genome coverage map of Bacteriophage MS2 constituted by all the mapped reads. The coverage of both the forward and reverse reads (top) and each strands separately (bottom) are shown. An almost equal population of both forward (44%) and reverse reads (56%) was found for Larifan, as represented by red and blue colours.

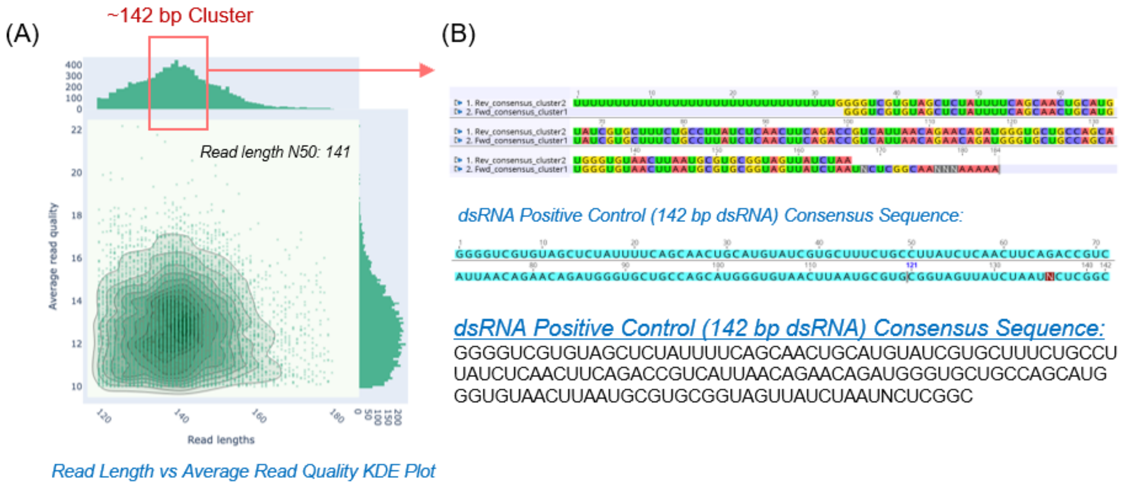


Figure S16. Nanopore sequencing of 142 bp dsRNA species (positive control dsRNA). (**A**) The read length vs. quality distribution is shown for the reads obtained from the 142 bp dsRNA sample-specific barcode. A clear abundance in the population around the 100-160 region can be seen, which largely represents the 142 bp dsRNA population. (**B**) The consensus mapping of forward and reverse strands, and the sequence of the RNA duplex species, is provided. This dsRNA species is fundamentally a synthetic sequence with no available BLASTn hit in the NCBI database.

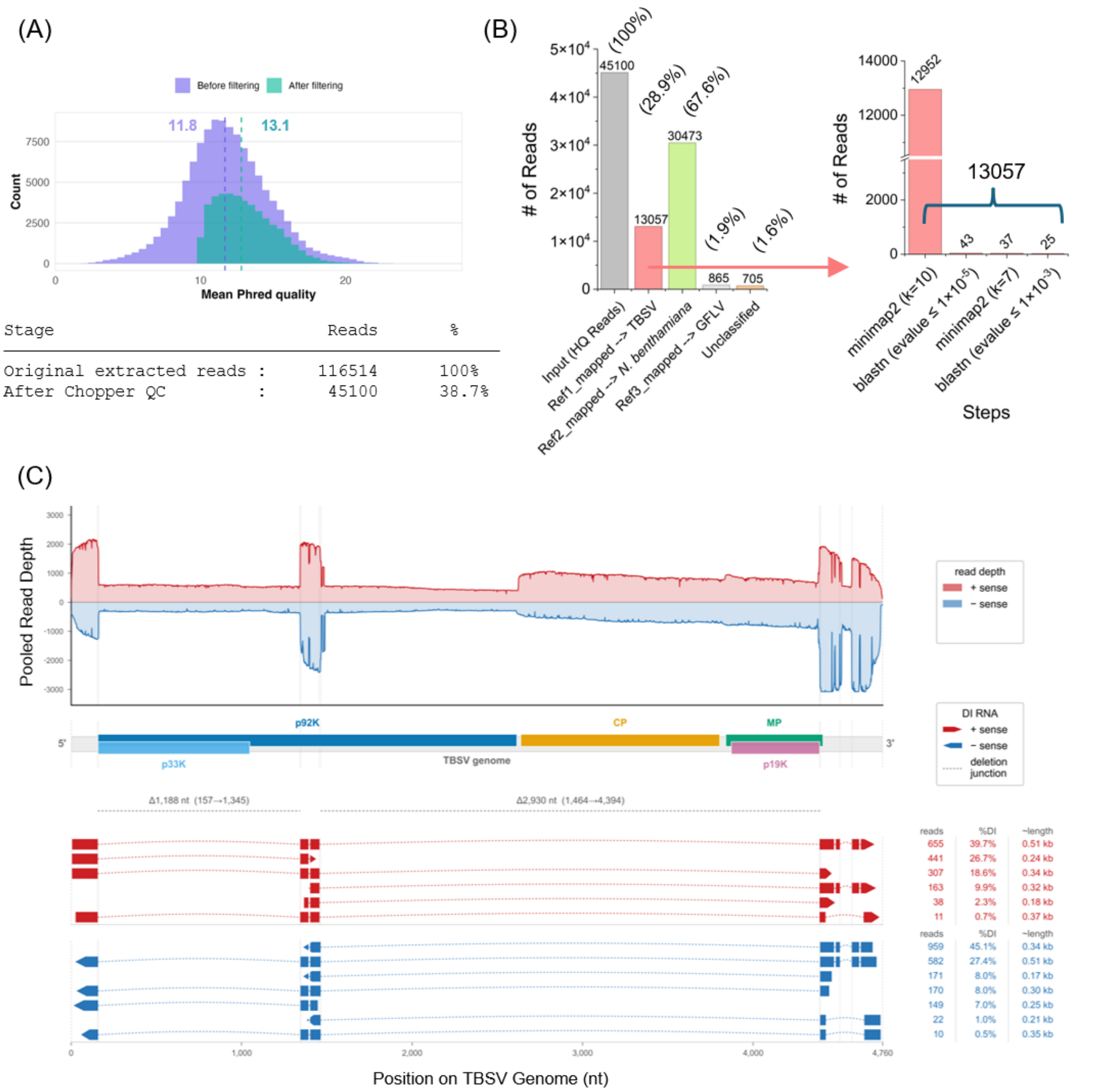


Figure S17. (**A**) Quality control filtering of all the TBSV-specific barcode (WDX_RTA_BC3 & 4) reads by the Chopper package (Q≥10). The mean Q score is shown for before and after filtering (11.8 and 13.1, respectively). Around 38% (45,100) of HQ reads are used for further mapping and classification analysis. (**B**) Mapping statistics of the TBSV-specific barcode reads to the virus and host genome. A total of 13,057 (~29%) reads map to the TBSV BS3Ng strain genome, while around 67% were host *N. benthamiana* reads. The target viral reads are obtained by a multi-step mapping approach using a combination of minimap2 and BLASTn tools (right) as described in the methods. (**C**) Analysis of the defective interfering (DI) RNAs found in the TBSV sample by the nanopore sequencing and overview of the BIRD-Seq pipeline. **(A)** Pooled read depth profile of the nanopore reads (top) mapped to TBSV genome revealing a non-uniform coverage with peaks at the 5'-end, internal ORF 92, and two 3'-terminal regions. Above x-axis: (+) sense strand (red); below x-axis: (-) sense strand (blue). Map of the TBSV genome showing the p33/p92 (replicase), CP, MP, and p19 cistrons are given below the read depth profile, and the dotted vertical lines show positions related to the deletion-junction. Mapping of the individual DI RNA species reads identified for the (+) sense (red) and (-) sense (blue) are provided at the bottom panel. The filled box regions represent the retained genomic segments, the connecting dotted lines reveal the deleted portions, and the end arrowheads show the strand orientation. The strand-specific DI RNA read length, number of reads, and percentage of population are given beside the map. The retained portions of the TBSV genome coincide with the prototypical tombusvirus DI RNAs signature.
